# Glioblastoma initiation, migration, and cell types are regulated by core bHLH transcription factors ASCL1 and OLIG2

**DOI:** 10.1101/2023.09.30.560206

**Authors:** Bianca L. Myers, Kathryn J. Brayer, Luis E. Paez-Beltran, Matthew S. Keith, Hideaki Suzuki, Jessie Newville, Rebekka H. Anderson, Yunee Lo, Conner M. Mertz, Rahul Kollipara, Mark D. Borromeo, Robert M. Bachoo, Jane E. Johnson, Tou Yia Vue

**Affiliations:** Department of Neurosciences, University of New Mexico Health Sciences Center, Albuquerque, New Mexico; University of New Mexico Comprehensive Cancer Center, Albuquerque, New Mexico; McDermott Center for Human Growth and Development, University of Texas Southwestern Medical Center, Dallas, Texas; Department of Neuroscience, University of Texas Southwestern Medical Center, Dallas, Texas; Department of Neurology and Neurotherapeutics, University of Texas Southwestern Medical Center, Dallas, Texas

**Author notes:** **Correspondence:** Tou Yia Vue.

## Abstract

Glioblastomas (GBMs) are highly aggressive, infiltrative, and heterogeneous brain tumors driven by complex driver mutations and glioma stem cells (GSCs). The neurodevelopmental transcription factors ASCL1 and OLIG2 are co-expressed in GBMs, but their role in regulating the heterogeneity and hierarchy of GBM tumor cells is unclear. Here, we show that oncogenic driver mutations lead to dysregulation of ASCL1 and OLIG2, which function redundantly to initiate brain tumor formation in a mouse model of GBM. Subsequently, the dynamic levels and reciprocal binding of ASCL1 and OLIG2 to each other and to downstream target genes then determine the cell types and degree of migration of tumor cells. Single-cell RNA sequencing (scRNA-seq) reveals that a high level of ASCL1 is key in defining GSCs by upregulating a collection of ribosomal protein, mitochondrial, neural stem cell (NSC), and cancer metastasis genes – all essential for sustaining the high proliferation, migration, and therapeutic resistance of GSCs.

## INTRODUCTION

Glioblastoma (GBM) is the most common and malignant form of glioma, with a median survival below 18 months and a five-year survival rate of less than 10%^1^. A major challenge in the treatment of GBM is that by the time of diagnosis, highly infiltrative tumor cells have already migrated across long distances on major white matter tracts and/or the microvasculature to invade surrounding brain regions^2^. Thus, despite the use of fluorescence guided surgery for maximal resection of tumor tissues followed by concurrent chemotherapy with temozolomide (TMZ) and conformal radiation^3–5^, recurrence is unavoidable since elimination of all residual cells is not possible. The limited efficacy of chemoradiation is also attributed to the heterogeneity and plasticity of GBM cells, whereby glioma stem cells (GSCs) identified via stem cell surface markers have been shown to be highly tumorigenic and resistant to TMZ^6–12^. Currently, the genetic mechanisms that regulate or determine GSCs or the hierarchy and heterogeneity of GBM tumors remain unclear.

Based on bulk RNA sequencing studies, GBMs are classified into three major subtypes – proneural, classical, and mesenchymal – each defined by a unique molecular signature^13–15^. However, single-cell RNA sequencing (scRNA-seq) and multi-location sampling of brain tumors reveal that individual tumors and tumor cells are comprised of multiple GBM subtypes, highlighting a high degree of inter- and intra-tumoral heterogeneity^16, 17^. More recently, single-cell lineage tracing combined with scRNA-seq demonstrates that GBM cells fluctuate between four transient cellular states resembling those of neural progenitor cells (NPCs), oligodendrocyte precursor cells (OPCs), astrocyte cells, and mesenchymal cells^18^. Although the relative frequencies of these cellular states and GBM subtypes are associated to some extent with specific genetic mutations/alterations or the tumor microenvironment^14, 15, 18, 19^, the transcriptomic plasticity of GBM cells is likely an inherent property independent of the types or combination of driver mutations.

A common feature of GBMs, including lower-grade gliomas, is the presence of multiple neurodevelopmental transcription factors that may be responsible for maintaining tumor cells in an undifferentiated, progenitor-like state^20–22^. Indeed, forced expression of several combinations of core transcription factors can reprogram differentiated glioma cells or transform immortalized astrocytes into tumor-propagating cells when transplanted orthotopically into brains of immunocompromised mice^20, 21^. These studies suggest that the combinatorial functions or differential levels of transcription factors are the direct mechanistic links between oncogenic driver mutations and downstream transcriptional programs to influence or determine the various cellular states and hierarchy of GBM cells. However, which transcription factors are responsible for which cellular states or cell types, and how the initiation, proliferation, and migration of GBM tumors are altered following perturbations of specific set of transcription factors *in vivo*, are unknown.

During development of the central nervous system (CNS), ASCL1 and OLIG2 are two influential transcription factors dynamically co-expressed in various multipotent NPCs, including in highly proliferative and migratory glial progenitor and precursor cells^23–30^. These two basic-helix-loop-helix (bHLH) transcription factors are co-expressed in an oscillatory manner with NOTCH signaling and HES proteins, and this oscillation is essential for balancing progenitor cell maintenance with cell fate specification of neuronal, oligodendroglial, and astroglial lineages^31, 32^. Similarly, ASCL1 and OLIG2 are highly co-expressed in GBM tumors, in part because loci of *OLIG1* and *OLIG2*, along with numerous NSC (*HES5, ID1, NFIX, SOX2, SOX4, ZEB1*) and glial lineage transcription factors (*ID3, NFIA, NFIB, NKX2-2, SOX8, SOX9, SOX10*), are major targets of ASCL1 binding^22, 33^. Developmentally, GBM cells follow a roadmap similar to that of the fetal brain, where ASCL1 marks the presence of highly proliferative, TMZ-resistant glial progenitor cancer cells at the apex of astroglial, oligodendroglial, neuronal, and mesenchymal cancer cells^34^. This similarity implies that, as seen during neurogenesis and gliogenesis, the dynamic levels of ASCL1 and OLIG2 directly underlie the process of gliomagenesis and the lineage composition of GBM tumors.

In this study, we show that ASCL1 and OLIG2 can physically and genetically interact to bind to similar sites in the genome of two separate patient derived orthotopic GBM xenografts (PDX-GBMs). By combining sophisticated transgenic gain- and loss-of-function strategies with an innovative CRISPR-Cas9 GBM mouse model in which brain tumors are induced and fluorescently labeled in the cortex, we demonstrate that ASCL1 and OLIG2 are two prominent drivers of all aspects of brain tumor initiation, proliferation, migration, and cell types. Notably, the co-expression but dynamic levels and inverse function of ASCL1 and OLIG2 are directly responsible for maintaining GSCs, glioma types, and the diverse cellular compositions of GBM tumors.

## RESULTS

### ASCL1 and OLIG2 share extensive overlap in binding in genome of orthotopic GBM xenografts

To understand the combinatorial function of ASCL1 and OLIG2 in GBMs, we performed ChIP-seq for OLIG2 for direct comparison with previously published ASCL1 ChIP-seq in two PDX-GBM lines^22^. We found that OLIG2 binds to 105,741 statistically significant sites (green circle, **Fig. 1A**) and overlapped with over 90% of the 13,457 ASCL1 binding peaks (green and blue circle overlap, **Fig. 1A,B**). *De novo* motif analyses showed significant enrichment of E-boxes with a “GC” core (CAGCGT) within ASCL1 and OLIG2 shared binding peaks, whereas E-boxes with a variable “GC/TA” core were observed within peaks for OLIG2 only. This difference in E-box binding preference and enrichment of different co-factor DNA motifs may explain the more promiscuous binding of OLIG2 compared to ASCL1 (**Fig. 1C**).

**Figure 1.**
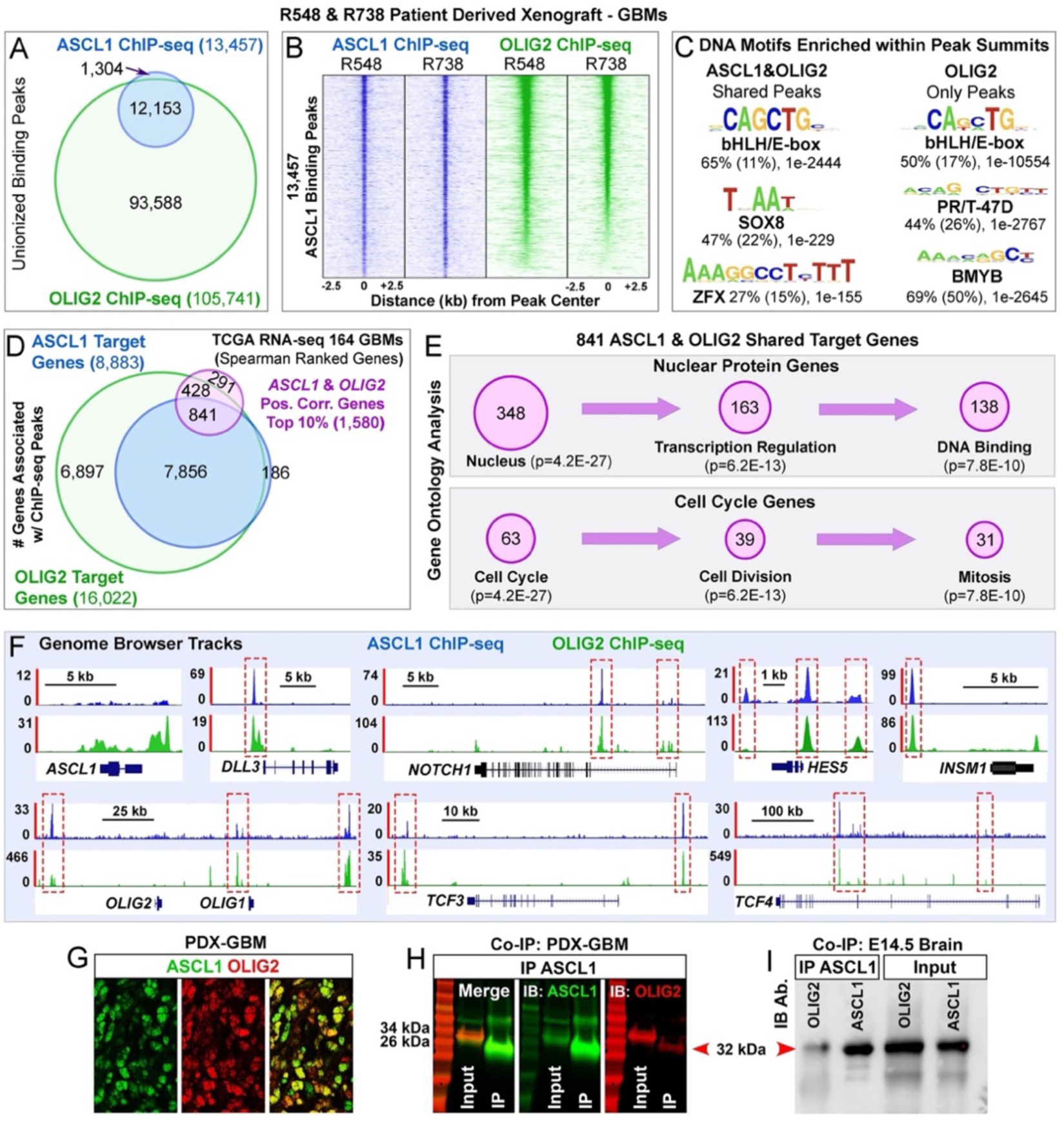
ASCL1 and OLIG2 physically interact and overlap in binding in genome of PDX-GBMs. (A) Venn diagram of ChIP-seq binding peaks for ASCL1 or OLIG2 from two different PDX-GBMs (R548 and R738). **(B)** Heatmap of ASCL1 and OLIG2 shared binding sites. **(C)** DNA sequence motifs that are enriched within ASCL1&OLIG2 or OLIG2 alone binding peaks. **(D)** Overlap of genes associated with ASCL1 or OLIG2 binding peaks intersecting with the top 10% of genes positively correlated with *ASCL1* and *OLIG2* expression in RNA-seq of 164 GBM samples from the TCGA. **(E)** Gene ontology analysis of 841 ASCL1 & OLIG2 shared targets demonstrating enrichment of molecular functions in the nucleus and cell cycle. **(F)** ASCL1 and OLIG2 binding peaks at their respective loci and known NOTCH (*DLL3, NOTCH1, HES5*), NSC (*INSM1*), and bHLH E-protein (*TCF3, TCF4*) targets. **(G)** Dynamic expression and colocalization of ASCL1 and OLIG2 in PDX-GBM. **(H,I)** Co-immunoprecipitation assay showing interactions between ASCL1 and OLIG2 in PDX-GBM and embryonic day (E)14.5 mouse brain. Scale bar: 25 μm.

To identify putative transcriptional targets of ASCL1 and OLIG2, we next used Genomic Regions Enrichment of Annotations Tool (GREAT)^35^ to associate the binding peaks of ASCL1 and OLIG2 with the nearest genes^22^. ASCL1 binding peaks were associated with 8,883 target genes, whereas OLIG2 binding peaks were associated with 16,022 target genes (**Supplementary Table 2 (TS2)**). Roughly 98% of the ASCL1 target genes were also targets of OLIG2 (**Fig. 1D**). Next, we applied Spearman rank-ordered correlation (>0.4) to identify the top genes positively correlated with both *ASCL1* and *OLIG2* in RNA-seq of 164 TCGA GBM samples (**TS3)**^13, 22^. This produced a total of 1,580 genes, 841 of which were direct targets and thus likely to be regulated by ASCL1 and OLIG2 binding (purple circle, **Fig. 1D**). Gene Ontology (GO) analysis of these 841 ASCL1 and OLIG2 shared targets revealed that about 40% (348) were associated with function within the nucleus, many of which play direct roles in transcription regulation and DNA binding. Additionally, about 7% (63) were associated with the cell cycle and are directly responsible for promoting cell division and mitosis (**Fig. 1E; TS4**). These functions are consistent with proposed roles for ASCL1 and OLIG2 as core regulators of tumor-propagating cells^21, 22, 34^.

We next investigated if ASCL1 and OLIG2 interact at both the genetic and protein levels, given their extensive shared binding as bHLH transcription factors. ChIP-seq analyses revealed the presence of ASCL1 and OLIG2 binding peaks at their respective *ASCL1* and *OLIG1/2* loci, and at known targets such as NOTCH signaling (*DLL1, DLL3*, *NOTCH1, HES1, HES5, HES6*) and NSC genes (*INSM1, ID1*). ASCL1 and OLIG2 binding peaks were also found at loci of *TCF3* and *TCF4* genes (**Fig. 1F**), which encode for bHLH E-protein DNA co-binding partners of ASCL1 and OLIG2. Co-immunoprecipitation assay demonstrated that the shared binding between ASCL1 and OLIG2 may be due to a direct protein-protein interaction since OLIG2 was successfully pulled down with an antibody specific for ASCL1 from PDX-GBM cells. This direct physical interaction was also found in NPCs of E14.5 mouse brains, which co-express high levels of ASCL1 and OLIG2 (**Fig. 1G-I**). Collectively, our ChIP-seq analyses suggest that ASCL1 and OLIG2 co-regulate a NSC transcriptional network essential to promote gliomagenesis.

### ASCL1 and OLIG2 play redundant roles in tumor initiation but inverse roles in regulating tumor cell migration and invasion in the brain of a GBM mouse model

We and others previously tested the requirement for ASCL1 or OLIG2 in two different GBM mouse models^22, 36^. Loss of either ASCL1 or OLIG2 had only modest effects on survival, likely due to the redundant binding between ASCL1 and OLIG2 (**Fig. 1**). Here, we hypothesized that the loss of both ASCL1 and OLIG2 should prevent or significantly compromise brain tumor induction or formation. To test this, we developed a CRISPR-Cas 9 transgenic GBM mouse model in which brain tumors are fluorescently labeled in wild-type immunocompetent mice. Specifically, plasmids expressing Cre-recombinase and Cas 9 + gRNAs^37^ targeting *Nf1, Pten*, and *Tp53* were injected and then electroporated into NPCs in the dorsal subventricular zone of the right lateral ventricle of Cre-dependent tdTomato (tdTOM) reporter mice (*R26R^T/T^*) at birth (P0) (**Fig. 2A; Fig. S1**). All electroporated mice consistently developed tdTOM+ tumors in the same location surrounding the right lateral ventricle in the striatum, corpus callosum, and cortex, and died between 2-3 months of age (**Fig. 2B-D,R**). The presence of tdTOM showed that tumors were aggressive and invasive, capable of migrating across the corpus callosum to the contralateral hemisphere, co-expressed high levels of both ASCL1 and OLIG2, and exhibited similar histopathological characteristics to that of GBMs (**Fig. 2D,E**; **Fig. 4D**). We then bred *Ascl1^Floxed^* and/or *Olig2^Floxed^* alleles into *R26^T/T^* mice to generate offspring to directly test the requirement of these two transcription factors specifically within tumor cells (**Table 1**), while non-tumor cells are normal.

**Figure 2.**
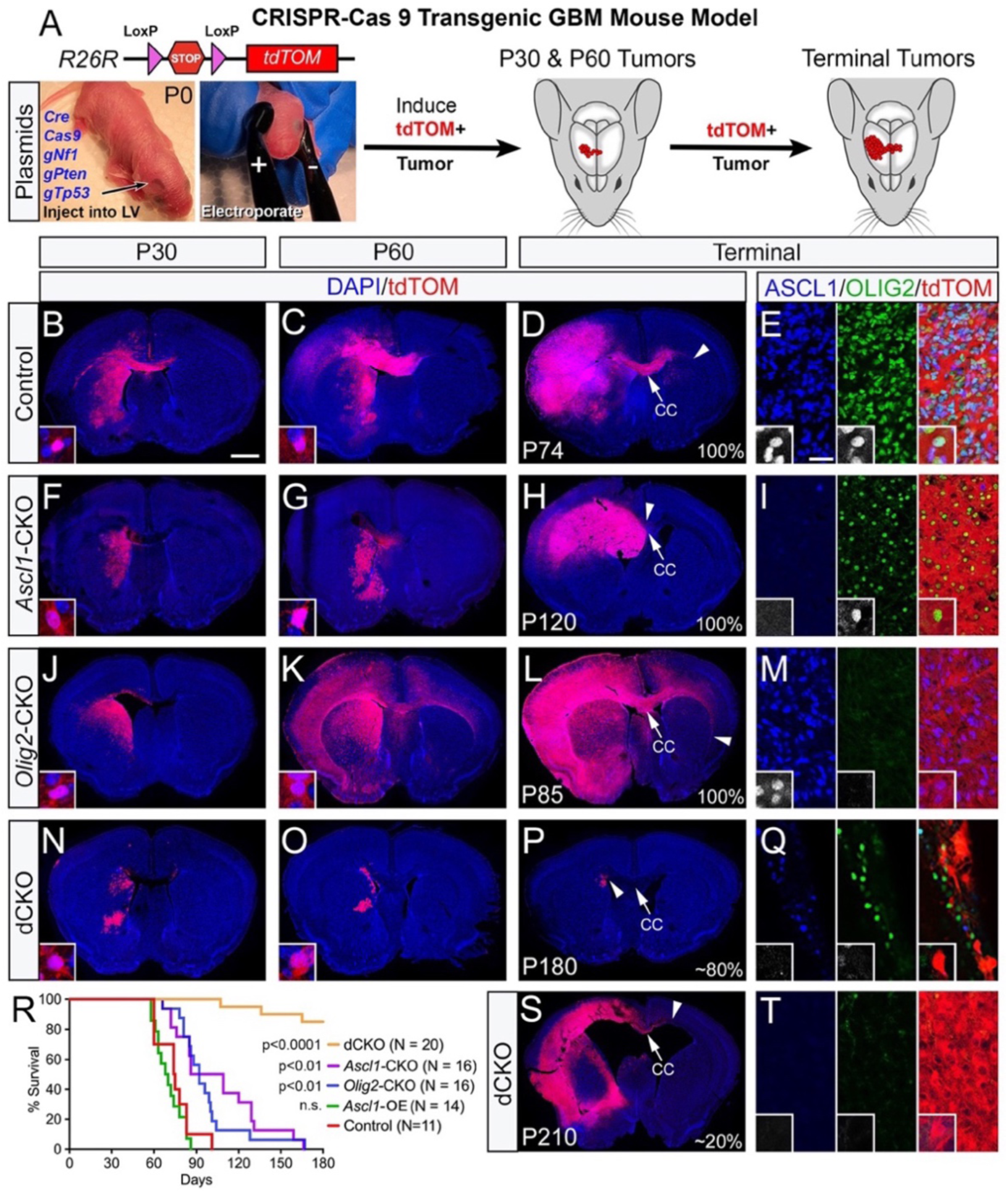
ASCL1 and OLIG2 are required for tumor formation but inversely regulate different aspects of tumor migration in GBM mouse model. (A) Induction of tdTOM+ brain tumors via electroporation of indicated Cre + CRISPR plasmids into neural progenitor cells in the right lateral ventricle of *R26R^T/T^* reporter mice at birth. (**B-Q,S,T)** Representative images of tdTOM+ tumors at P30, P60, or terminal stage in control (B-E), *Ascl1*-CKO (F-I), *Olig2*-CKO (J-M), and double CKO (N-Q,S,T) mice. Arrows indicate midline and arrowhead marks the distance of migration of tdTOM+ tumor cells on the contralateral corpus callosum. ASCL1 and OLIG2 are highly co-expressed in control tdTOM+ tumor cells, but absent in the single or double CKO tdTOM+ tumors. **(R)** Survival curves of each group of tumor mice showing statistical significance (Kaplan-Meier) between control versus experimental groups. Scale bars: 1 mm for whole brain sections; 25 μm for panels E,I,M,Q,T, and 12.5 μm for all insets.

**Table 1.**
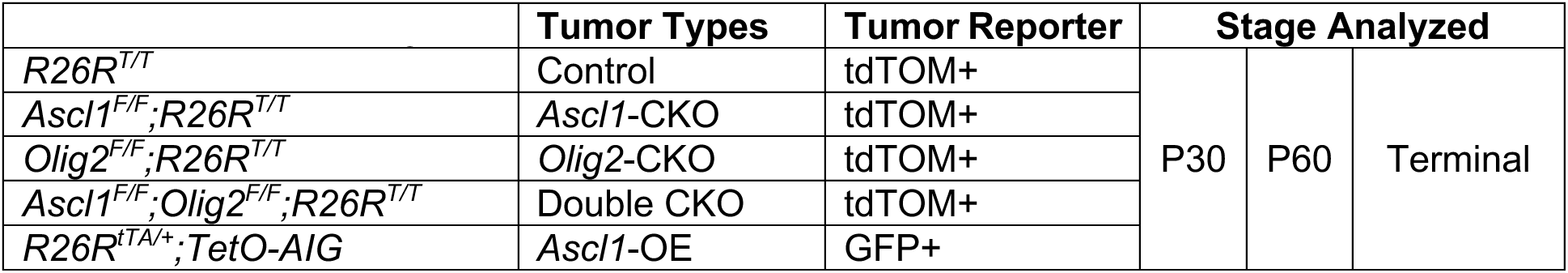
Mouse Genotypes.

As in our previous study^22^, *Ascl1*-CKO resulted in tumor formation and a longer median survival time (98 days versus 75 days for control mice) (**Fig. 2R**), which was reflected by smaller tumors at P30 and P60 (**Fig. 2F,G**). However, there was a reduction in migration along the corpus callosum to the contralateral hemisphere, even at terminal stages, although OLIG2 was still present (**Fig. 2H,I**). Overall, the distance of tdTOM+ tumor cell migration along the contralateral corpus callosum was decreased by 50% compared to control tumors, indicating that ASCL1 is a positive regulator of tumor cell migration (**Fig. 3I**).

Similar to *Ascl1*-CKO, *Olig2*-CKO resulted in efficient tumor formation with a median survival of 92 days (**Fig. 2R**), likely because ASCL1 was still present in these tumors (**Fig. 2M**). However, unlike *Ascl1*-CKO tumors, *Olig2*-CKO tumors were highly diffused, indicating that OLIG2 is a negative regulator of tumor cell migration. Indeed, from P60 to terminal stages, *Olig2*-CKO tumors migrated to occupy almost the entire contralateral corpus callosum and cortex (**Fig. 2J-L**), and were accompanied by a 2-to 4-fold increase in migration distance on the contralateral corpus callosum and a significant decrease in tumor cell density on the ipsilateral hemisphere compared to control and *Ascl1*-CKO tumors (**Fig. 3H,I**).

As predicted, double CKO (dCKO) prevented tumor formation in approximately 80% of mice (N=16/20, 4 litters) (**Fig. 2P,Q**), while 20% developed slow-growing tumors that were not lethal until 4-6 months of age (**Fig. 2R-T**), consistent with the redundant function of ASCL1 and OLIG2. Indeed, compared to control or single CKO mice, there were far fewer dCKO tdTOM+ cells surrounding the right ventricle in the striatum or corpus callosum at P30 or P60 (**Fig. 2N,O**). Furthermore, the size and number of these dCKO tdTOM+ cells decreased with age by 6 months and failed to migrate onto the corpus callosum or the overlying cortex in the non-tumor mice (**Fig. 2N-P**). A few of the dCKO lethal tumors that developed in 20% of the mice did show extensive ipsilateral migration into the cortical grey matter and some contralateral migration on the corpus callosum (**Fig. 2S**), but this migration was not as drastic compared to control or *Olig2*-CKO tumors (**Fig. 2D,L**).

We also analyzed whether sex had an effect on tumor development and found that there was no statistical difference in survival between male and female mice for the various tumor types (**Fig. S1G,H**). Taken together, our findings imply that ASCL1 and OLIG2 function redundantly downstream of driver mutations to transform affected NPCs into proliferating tumor cells, but these transcription factors regulate opposing aspects of tumor cell migration.

### High levels of ASCL1 promote a highly migratory and diffuse glioma phenotype

The fact that ASCL1 and OLIG2 can physically and genetically interact (**Fig. 1F-I**) suggests that OLIG2 may directly repress ASCL1’s ability to promote tumor migration through these interactions. We hypothesized that if ASCL1 is directly required to promote tumor migration, then increasing ASCL1 levels should overcome the repression by OLIG2, resulting in a highly migratory and diffuse tumor phenotype similar to *Olig2*-CKO tumors. To test this, we induced tumors in transgenic mice carrying dual alleles of a Cre-dependent tetracycline transactivator and a TetO-promoter driving expression of *Ascl1-ires-GFP* (**Fig. 3A**)^38^, hereafter referred to as *Ascl1*-overexpression (OE) tumors. Analysis of cellular immunofluorescent intensity confirmed that ASCL1 was elevated about 2-fold in *Ascl1*-OE GFP+ tumor cells compared to control tumor cells (**Fig. 3F**).

**Figure 3.**
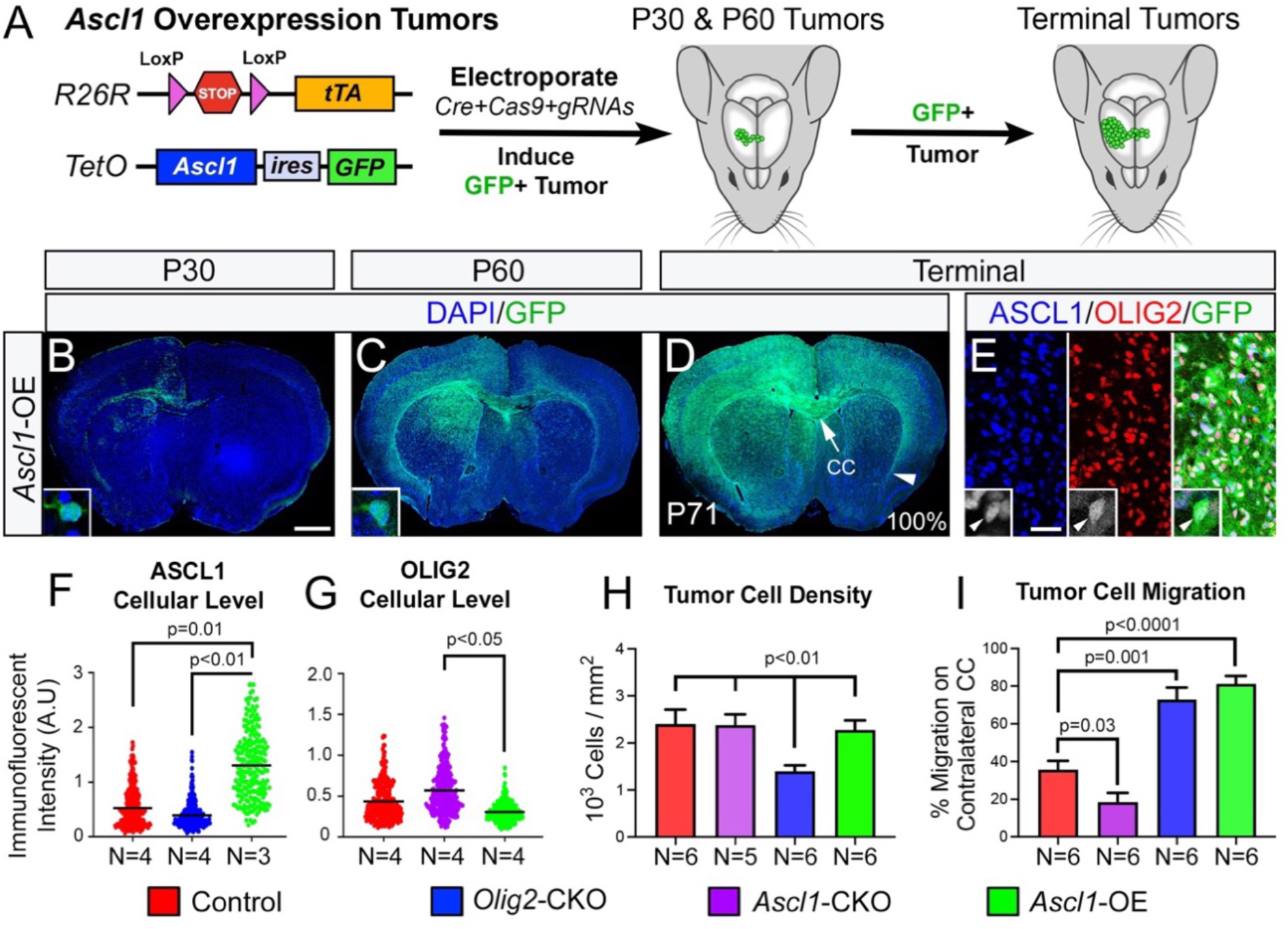
ASCL1 overexpression (OE) promotes tumor cell migration. (A) *Ascl1-*OE GFP+ tumor model. **(B-E)** Representative images of P30, P60, and terminal stage tumors highlighting extensive migration and co-expression of ASCL1 and OLIG2 in GFP+ tumor cells. **(F,G)** Scatter plot of immunofluorescent intensity of ASCL1 or OLIG2 within individual DAPI+ nuclei of tumor cells of genotypes indicated. (**H,I**) Quantification of the density of DAPI+ tumor cells in tumor bulk, and distance of migration of reporter+ tumor cells on contralateral corpus callosum normalized to the total length of the contralateral corpus callosum. Data shown as mean ± SEM. Statistical significance are determined by comparing the means of tumor types using unpaired t-tests. Scale bars: 1 mm for panels B,C,and D; 25 μm for panel E; and 12.5 μm for all insets.

As expected, GFP+ tumor cells had migrated extensively on the ipsilateral corpus callosum and cortex at P30 (**Fig. 3B**). By P60 to terminal stages, GFP+ tumor cells had infiltrated the striatum and cortex of both ipsilateral and contralateral hemispheres (**Fig. 3C,D**), similar to *Olig2*-CKO tumors. Migration distance of *Ascl1*-OE GFP+ tumor cells on the contralateral corpus callosum was similar to that of *Olig2*-CKO tumors (**Fig. 3H**). However, the levels of ASCL1 in *Olig2*-CKO tumor cells, though not dissimilar to control, were much lower compared to *Ascl1*-OE tumor cells (**Fig. 3F**). Conversely, the levels of OLIG2 were higher in *Ascl1*-CKO compared to control but were significantly reduced in *Ascl1*-OE tumor cells (**Fig. 3G**). These findings highlight the general observation that the cellular levels of ASCL1 and OLIG2 are highly dynamic but inversely proportional to each other within tumor cells. It is possible that the degree of this inverse levels of ASCL1 relative to OLIG2 that directly determine the degree of migration or diffusion of glioma tumors.

### ASCL1 and OLIG2 also play inverse roles in regulating glioma tumor types and cell types

Based on their developmental roles, we hypothesized that altering the levels of ASCL1 and/or OLIG2 should influence the cell types in tumors of the GBM mouse model. Within control tumors, tdTOM colocalized extensively with either GFAP or SOX10, but rarely with both within the same cells (**Fig. 4A-C**). This indicates that these tumors are “mixed gliomas” of astroglial or oligodendroglial lineages. While *Ascl1*-CKO tumors also had an abundance of GFAP+ and SOX10+ cells within the tumor bulk, most tdTOM+ tumor cells were SOX10+ and very few were GFAP+ (**Fig. 4E-G**). This suggests that *Ascl1*-CKO tumors are predominantly “oligodendrogliomas” and most GFAP+ cells are likely normal or reactive astrocytes. Conversely, *Olig2*-CKO tumors are “astrocytomas” dominated by extremely high levels of GFAP (**Fig. 4I-K**). While SOX10+ cells are present within *Olig2*-CKO tumors, these cells were not tdTOM+ (**Fig. 4K**), and therefore are non-tumor oligodendrocyte lineage. Similarly, tdTOM+ cells of dCKO tumors are predominantly GFAP+ and SOX10 is absent (**Fig. 4J-L**), indicating that they are also astrocyte-like cells. However, the levels of GFAP in the few dCKO tumors (**Fig. 4M**) are lower than those of *Olig2*-CKO tumors (**Fig. 4I**). Given that *GFAP* is a target of both ASCL1 and OLIG2 binding (**Fig. 7B**), the extremely high levels of GFAP in *Olig2*-CKO tumors are likely due to the presence of ASCL1, and not just the lack of repression of astrocyte fate by the loss of OLIG2, which also occurs in dCKO tumors. Finally, GFP+ cells of *Ascl1*-OE tumors, which highly co-express OLIG2 (**Fig. 3E**), were mostly either GFAP+ or SOX10+ (**Fig. 4Q-S**), and thus were also “mixed gliomas”. However, the levels of GFAP were noticeably higher in *Ascl1*-OE tumors (**Fig. 4Q**) compared to control tumors (**Fig. 4A**), which could be due to the imbalance of high levels of ASCL1 to lower levels of OLIG2 in these tumor cells (**Fig. 3F,G**). The glioma types of these tumors were also confirmed by H&E staining revealing the predominance of pink GFAP fibers (arrows) or round “fried egg” cells with perinuclear halo (arrowhead), which are characteristics of astrocytoma and oligodendroglioma, respectively (**Fig. 4D,H,L,P,T**).

**Figure 4.**
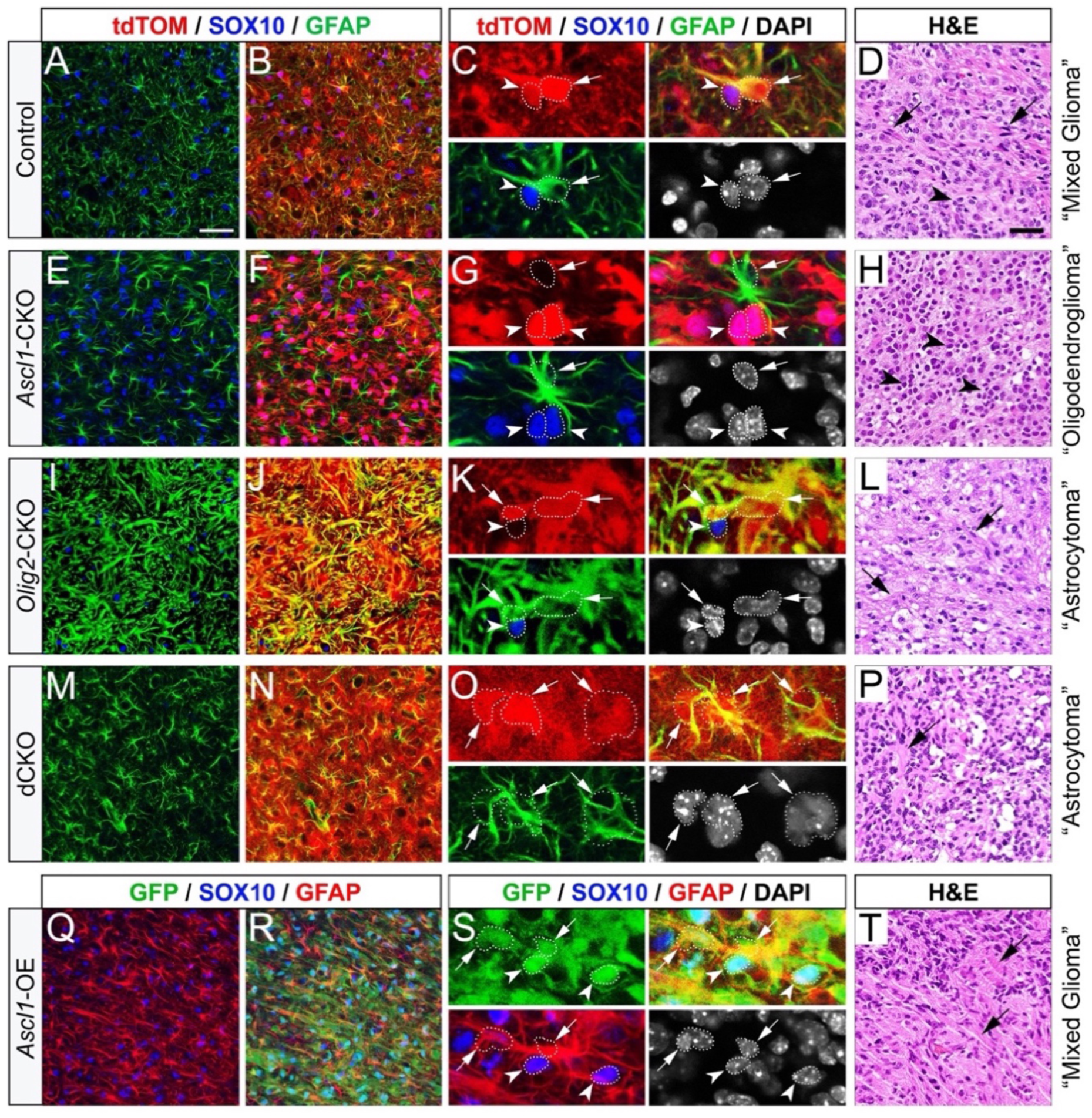
ASCL1 and OLIG2 inversely regulate glioma tumor cell types. Representative H&E staining and immunofluorescent images of terminal control (**A-D**), *Ascl1*-CKO (**E-H**), *Olig2*-CKO (**I-L**), dCKO (**M-P**), and *Ascl1*-OE (**Q-T**) tumors. High magnification showing differences in co-localization, or lack thereof, of GFAP (arrow) or SOX10 (arrowhead) in reporter+ tumor cells. Dotted line delineates DAPI+ cell nuclei that overlap with tumor reporter and/or cell type markers. Arrows and arrowheads in H&E panels indicate present of GFAP+ pink fibers and “fried-egg” cell shapes characteristics of astrocytoma and oligodendroglioma, respectively. Scale bars: 12.5 μm for panels C,G,K,O,S, and 50 μm for all other panels.

We also analyzed expression of SOX2 and PDGFRA markers for all tumor types. Normally, SOX2 is expressed at high levels in NSCs/NPCs but at lower levels in astrocytes and OPCs^39, 40^, whereas PDGFRA is specific to OPCs^41^. We observed that tumor cells, irrespective of genotypes, were SOX2+ (**Fig. S2**). In contrast, PDGFRA was strongly detected in OLIG2+ tumors (control, *Ascl1*-CKO, *Ascl1*-OE) (**Fig. S2B,F,R**), but was absent or greatly reduced in tumors without OLIG2 (*Olig2*-CKO and dCKO) (**Fig. S2J,N**).

Overall, our lineage analysis demonstrates that the glial cell fates of tumor cells derived from NPCs in the subventricular zone reflect that seen during gliogenesis. More importantly, all tumor cells were SOX2+ progenitor-like cells, with ASCL1 promoting an astroglial-like fate and OLIG2 promoting an oligodendroglial-like fate.

### ASCL1 and OLIG2 positively regulate each other’s expression and function redundantly to promote tumor cell proliferation

We next sought to determine if expression of ASCL1 or OLIG2 was altered at both early and terminal stages of the various tumor types. Within control tumors, we found that 74% of tdTOM+ tumor cells were OLIG2+ at P30, but was decreased to 61% at terminal stages (**Fig. S3A,B,K**). In contrast, 35% of control tdTOM+ tumor cells were ASCL1+ at P30, which then increased to 54% at terminal stages (**Fig. S3E,F,L**). Interestingly, *Ascl1*-CKO or *Olig2*-CKO significantly reduced the expression of the other transcription factor at P30 compared to control, but this reduction was not observed in terminal tumors (**Fig. S3K,L**). In contrast, *Ascl1*-OE significantly increased the number of OLIG2+ tumor cells in both P30 and terminal tumors compared to control (**Fig. S3K**). Thus, despite their inverse functions, these findings indicate a positive regulatory relationship between ASCL1 and OLIG2.

Since ASCL1 and OLIG2 are able to directly bind to cell cycle genes (**Fig. 1F**), we next investigated how alteration of the levels of ASCL1 and/or OLIG2 affect tumor cell proliferation. We injected tumor mice with EdU, a thymidine analogue that is incorporated into replicating cells during S-phase, for 2 hours prior to tumor harvest at P30 or terminal stages. We then analyzed regions of brain tumors with the highest density of EdU+ cells for all tumor types. Within control tumors, 5% of tdTOM+ tumor cells were EdU+ at P30, and this was increased to 9% at terminal stages (**Fig. S4A,B,K,L**). Interestingly, neither *Ascl1*-CKO nor *Olig2*-CKO had a significant effect on tumor cell proliferation at P30 or terminal stages (**Fig. S4C-F,K,L**). In contrast, dCKO showed no incorporation of EdU within tdTOM+ cells at P30, or even in the terminal stage dCKO tumors (**Fig. S4G,H**), highlighting a complete lack of cell proliferation in the absence of both ASCL1 and OLIG2. Conversely, *Ascl1*-OE GFP+ tumor cells had a significantly higher percentage of EdU+ cells at P30, and to a lesser extent at terminal stage compared to control or single CKO tumors (**Fig. S4I-L**). This increased proliferation of *Ascl1*-OE tumor cells is likely responsible for high tumor cell density compared to *Olig2*-CKO tumors despite the similarity in migratory phenotypes (**Fig. 4H**).

Taken together, these findings revealed that ASCL1 and OLIG2 are differentially dysregulated by the loss of *Nf1, Pten*, and *Tp53* at early stages (P30), but these two transcription factors are able to reciprocally and positively regulate each other’s expression to redundantly promote tumor formation and progression.

### Tumor cells of the GBM mouse model are highly heterogeneous and contain all GBM subtypes

To determine if tumors of the GBM mouse model exhibit a high degree of heterogeneity at the single cell level similar to humans, and how ASCL1 is directly responsible for regulating the transcriptome of tumor cells, we performed scRNA-seq analyses on control and *Ascl1*-OE tumors (**Fig. 5A**). We chose these two tumor types because they both co-express ASCL1 and OLIG2, exhibited similar “mixed glioma” cell types, but show completely different migratory behavior. To avoid possible confounding transcriptomes from non-tumor cells (i.e. reactive astrocytes, microglia/macrophages), we performed scRNA-seq only on FAC-sorted tdTOM+ cells of control tumors (N=3, ∼18,000 cells) or GFP+ cells of *Ascl1*-OE tumors (N=3, ∼25,000 cells) and detected similar numbers of genes per cell for both tumor types (**Fig. 5B,C**). Using Cell Ranger pipeline analyses, transcripts of tumor cells were normalized, integrated, and then grouped into cell clusters for visualization using Uniformed Manifold Approximation and Projection (UMAP) analyses. We detected 28 transcriptionally diverse cell clusters across both tumor types (**Fig. 5D,E; Fig. S5A-H**), but some clusters were significantly more enriched in control or *Ascl1*-OE tumors (circles and arrows, **Fig. 5D-F**). Gene expression analyses confirmed that *Ascl1*, along with the canonical NOTCH signaling targets *Dll3* and *Hes5* (**Fig. 5G-I**) and the bHLH E-protein co-binding partners, *Tcf4* and *Tcf12* (**Fig. S5I,J**), were significantly elevated in *Ascl1*-OE compared to control tumor cells.

**Figure 5.**
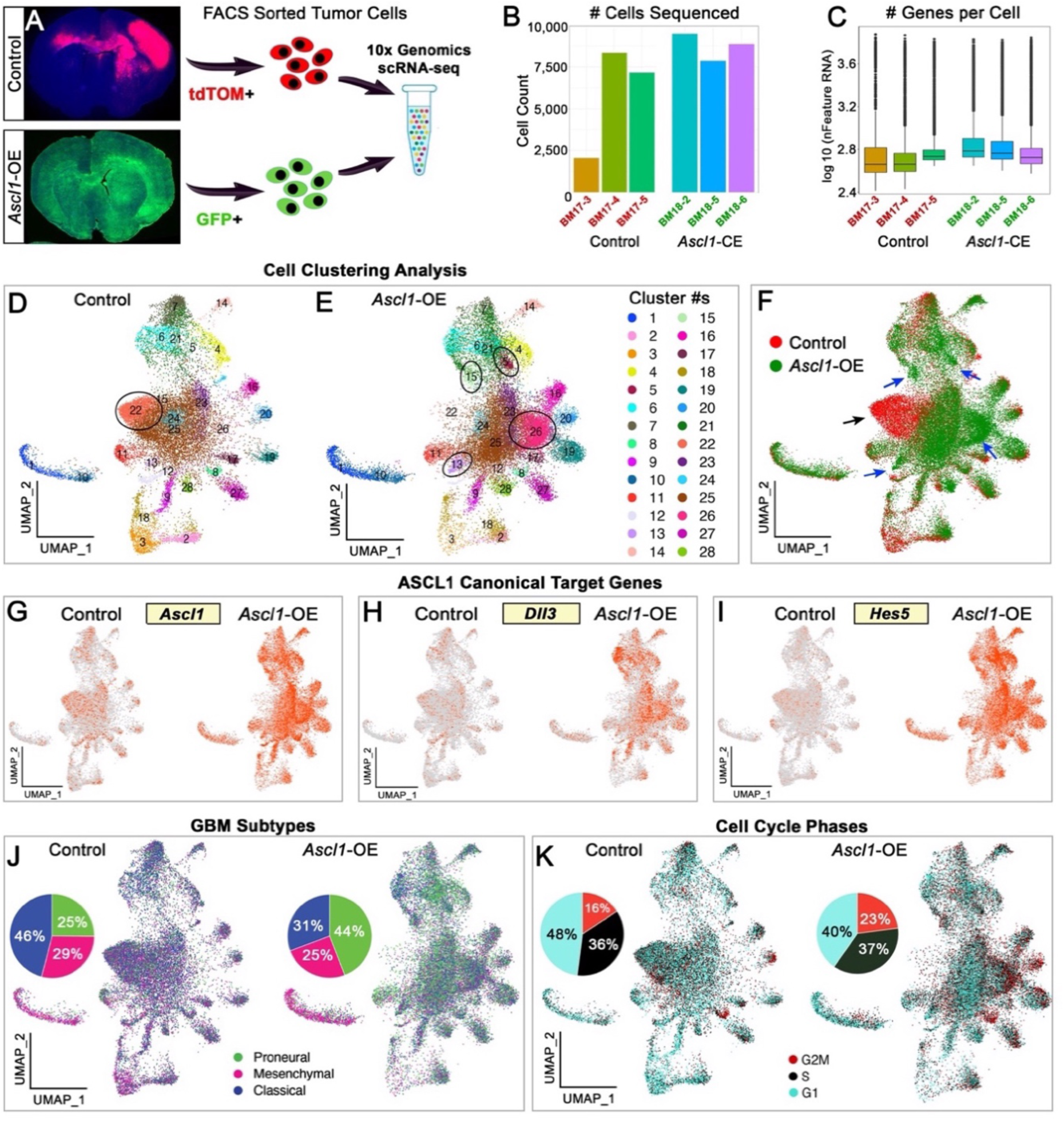
Single-cell RNA-seq reveals high degrees of transcriptional diversity for control and *Ascl1*-OE tumor cells. (A) Workflow of FACS-sorted tdTOM+ or GFP+ tumor cells for scRNA-seq. **(B-C)** Number of cells per tumor and genes per cell sequenced for control (N=3) and *Ascl1*-OE (N=3) tumors. **(D-F)** UMAP visualization of unsupervised clustering of control and *Ascl1*-OE tumor cells yielded 28 different cell clusters. Cell clusters enriched with control (black arrow) or *Ascl1*-CE tumors (blue arrows) are indicated (ovals). (**G-H**) Gene expression confirming increased levels of *Ascl1, Dll3,* and *Hes5* in *Ascl1*-OE tumor cells compared to controls. (**J,K**) GBM subtypes and cell phase analyses showing enrichment of proneural and decreased of classical GBM subtypes, and increased proportions of cells in G2M phase with *Ascl1*-OE.

We next examined if the three GBM subtypes — proneural, neural, and mesenchymal — were represented within the 28 cell clusters and how they were affected by the increased levels of ASCL1 in *Ascl1*-OE tumors compared to control tumors. GBM subtype identity was assigned to all tumor cells using the top 50 signature genes for each subtype^14, 15^. All three GBM subtypes were represented within cells from both tumor models, but the predominance of these subtypes differed in *Ascl1*-OE tumors, in which the proneural subtype was increased at the expense of the classical subtype (**Fig. 5J**). Cells in G2M phase were also increased in *Ascl1*-OE tumors (23%) compared to control tumors (16%) (**Fig. 5K**), supporting our previous findings that ASCL1 promotes tumor cell proliferation (**Fig. S3K,L**)^22^.

### High levels of ASCL1 promote NSC/astrocyte-like cell types at the expense of OPC/oligodendrocyte-like cell types in brain tumors

We next analyzed how the lineage development and lineage composition of tumor cells were altered in *Ascl1*-OE versus control tumors. We assigned a predicted cell type based on gene signatures (**TS5**) derived from previously reported cell type-specific bulk RNA-seq or scRNA-seq data sets of CNS and non-CNS cells in the mouse brain. These included quiescent and active NSCs from embryonic mouse brains^42–44^, oligodendrocyte lineage developmental cell states (OPC, NFOL, MOL) characterized in juvenile and adult CNS^45–47^, and astrocytes, neurons, and non-CNS (microglia, endothelial, and pericytes) cells acutely purified from brains of juvenile mice^45–47^.

In both control and *Ascl1*-OE tumors, approximately 80% of the tumor cells were assigned a CNS cell type (OPC, NFOL, MOL, quiescent or active NSC, astrocyte, or neuron), while ∼20% were assigned a non-CNS cell type (microglia, endothelial, pericyte). Most cell types were intermixed within the 28 UMAP cell clusters into three major cluster (MC) groups, which we termed MC1 through MC3 (**Fig. 6**). MC1 was comprised of about 80% (22/28) of cell clusters and all 10 assigned cell types, although at varying proportions. MC2 was comprised of 4 cell clusters (#2,3,9,18) and contained predominantly NFOL and MOL assigned cell types (**Fig. 6A,D**). MC3 was the smallest and most segregated cluster, comprised mainly of microglial cell types (**Fig. 6C,F**), also identified as mesenchymal GBM subtypes (**Fig. 5J**).

**Figure 6.**
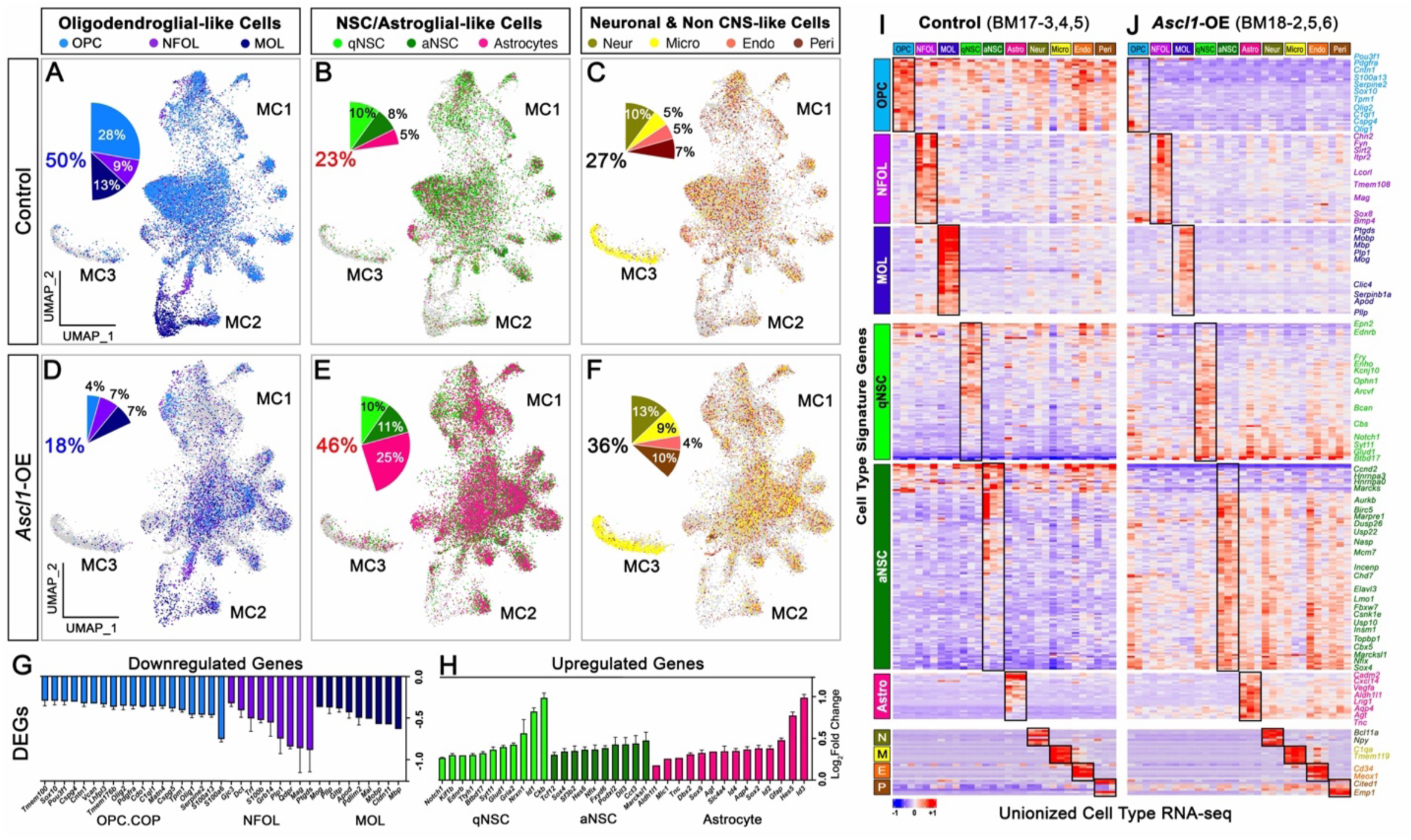
High levels of ASCL1 promote NSC/astroglial-like cells and suppress OPC/oligodendroglial-like cells in mouse GBM tumors. (A-F) UMAPs demonstrating the proportion and distribution of 10 assigned cell types in control (A-C) and *Ascl1*-OE (D-F) tumors based on cell-type specific gene signatures. **(G-H)** Differential gene expression confirms downregulation of oligodendrocyte lineage-specific genes (G) and upregulation of NSC/astrocyte-specific genes (H) in *Ascl1*-OE tumor cells. (**I,J)** Heatmap of unionized transcriptome for each assigned cell types into pseudo-bulk RNA-seq triplicates for control and *Ascl1*-OE tumors showing specificity of signature genes to their respective cell types (black rectangles). All cell types of control tumors expressed some level of OPC and NSC signature genes, which were the most drastically downregulated or upregulated in *Ascl1*-OE tumors, respectively.

About 50% of control tumor cells were of oligodendrocyte lineages (28% OPC, 9% NFOL, 13% MOL) (**Fig. 6A**), whereas 23% were of NSC (10% qNSC, 8% aNSC) and astrocyte (5%) lineages (**Fig. 6B**); the remaining were neuronal (10%) and non-CNS (17%) cell types (**Fig. 6C**). In contrast, *Ascl1*-OE tumors showed a 5-fold increase in astrocyte cell types (25%) versus oligodendrocyte lineage cell types, in which OPC (4%) and MOL (7%) were decreased by 7-fold and 2-fold, respectively, compared to controls (**Fig. 6D,E**). Proportions of neuronal and non-CNS assigned cell types were not significantly affected by *Ascl1*-OE (**Fig. 6F**). The switch in assigned cell types in *Ascl1*-OE tumors was accompanied by specific downregulation of OPC, NFOL, and MOL signature genes and upregulation of NSC and astrocyte signature genes compared to control tumors (Log**2**FC **≥ ±**0.25) (**Fig. 6G,H**).

We then combined the transcriptomes of each assigned cell type into pseudo-bulk RNA-seq analyses to better characterize the specificity of the signature genes and how their expression was altered at the population/cell type level in *Ascl1*-OE tumors compared to controls. This resulted in triplicates of unionized transcriptome for each cell type from the three control (BM17-3,4,5) and three *Ascl1*-OE (BM18-2,5,6) tumors (**Fig. 6I,J**). Heatmap analyses demonstrated that, as expected, expression of signature genes was largely specific to their respective assigned cell types (black rectangles, **Fig. 6I,J**), but many OPC and a subset of quiescent and active NSC genes were also expressed across all other assigned cell types, indicating that the more mature tumor cell types are derived from the OPC and NSC tumor cells. At this global level, oligodendrocyte lineage (OPC, NFOL, MOL) signature genes, especially for OPC, were significantly downregulated in *Ascl1*-OE tumors across all cell types. On the other hand, many NSC and astrocyte signature genes, especially those expressed at low levels in control tumors, were markedly upregulated in all cell types of *Ascl1*-OE tumors, whereas most neuronal and non-CNS cell type signature genes were largely unaffected. This pattern of differential regulation was also observed when displayed qualitatively across UMAP cell clusters, even though many of the OPC/oligodendroglial (*Olig2, Sox10, Pdgfra, Pou3f1, S100a6, Mag, Mbp*), NSC/astroglial (*Sox2, Sox4, Gfap, Id1, Id3, Aldh1l1, Aqp4*), neuronal (Npy, Stmn2) and non-CNS (*Irf8, Trem2, Bsg, Meox1, C4b, Emp1*) cell type signature genes were targets of ASCL1 binding (**Fig. 7; TS2**).

**Figure 7.**
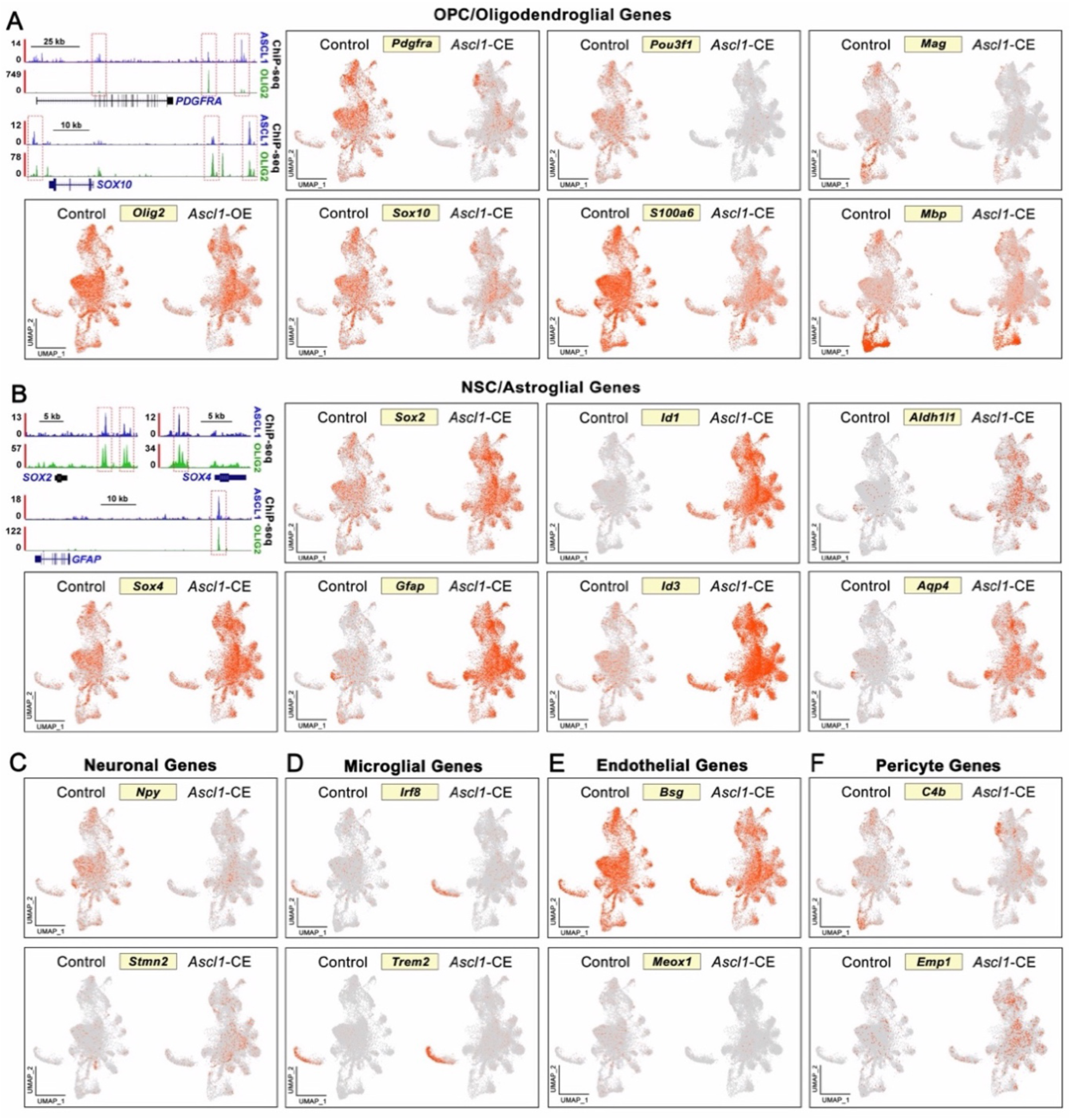
ASCL1 and OLIG2 shared target cell type specific genes are differentially expressed in control and *Ascl1*-OE tumors. (A,B) ChIP-seq tracks demonstrating shared binding of ASCL1 and OLIG2 at cis-regulatory cites of OPC/oligodendroglial (A) and NSC/astroglial (B) target genes and UMAP showing differential expression in control and *Ascl1*- OE cell clusters. **(C-F)** UMAPs showing differential expression of neuronal, microglial, endothelial, and pericyte lineage-specific genes. Note, all genes except for *S100a6* and *Mag* are targets of ASCL1 and OLIG2.

### Ribosomal protein, mitochondrial, and cancer metastasis genes are upregulated in NSC/astrocyte-like cells of *Ascl1*-OE tumors

To identify novel genes (whether directly or indirectly regulated by ASCL1) that may contribute to the highly invasive phenotype of *Ascl1*-OE tumors, we first identified differentially expressed genes (DEGs, Log**2**FC **;: ±**0.25) up- or downregulated in at least 10% (3/28) of cell clusters versus control tumor cells. We found that 275 genes were upregulated, whereas 109 genes were downregulated, and about 40% of each of these DEGs were identified as targets of ASCL1 binding based on ChIP-seq (**Fig. 8A**). The top genes and biological processes enriched within the 275 upregulated genes were ribosomal protein and translation elongation factor encoding genes essential for sustaining the necessary protein synthesis required for highly proliferative and migratory cancer cells^48^. In fact, an astounding 71 of 80 ribosomal protein small/large (*Rps/Rpl*) subunit genes and 4 of 5 translation elongation factor genes were significantly upregulated. The next most enriched gene set was related to oxidative phosphorylation, specifically mitochondrial-encoded (*mt-Co1, mt-Co2, mt-Co3*) and nuclear-encoded (*Cox4i1, Cox5a, Cox5b, Cox6a, Cox8a*) cytochrome oxidase subunit genes essential for Complex IV of the electron transport chain (**Fig. 8B,D,E**). The upregulation of these translational and ATP synthesis related genes was observed across all assigned cell types, but was highest within the microglial, endothelial, and pericyte tumor cell types (**Fig. 8C,D**).

**Figure 8.**
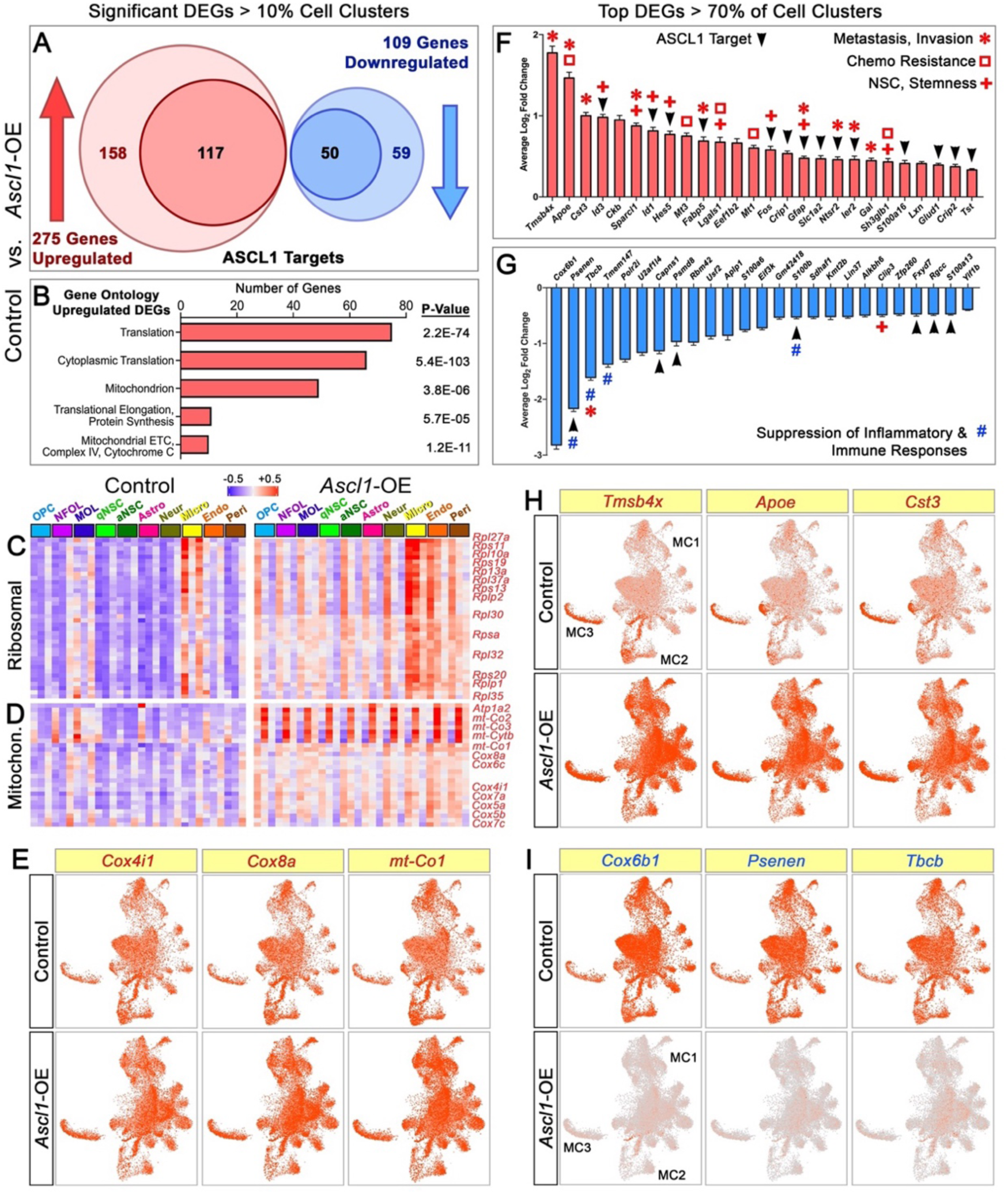
Ribosomal protein, mitochondrial, and genes important for NSC, cancer metastasis, and therapeutic resistance are upregulated in *Ascl1*-OE tumor cells. **(A)** Differentially expressed genes (DEGs) significantly up- or downregulated in more than 10% of cell clusters, many of which are targets of ASCL1. **(B)** Gene ontology analysis of upregulated DEGs showing enrichment of genes important for protein synthesis and mitochondrial function. **(C-D)** Heatmap from pseudo-bulk cell type RNA-seq demonstrating upregulation of ribosomal (C) and mitochondrial (D) genes in *Ascl1*-OE tumors. **(E)** UMAPs showing specific upregulation of cytochrome C oxidase subunit genes. **(F-G)** Top genes up- (F) or downregulated (G) in >70% of cell clusters, with delineation as ASCL1 targets and known functions in cancer or GBMs. **(H-I)** UMAP gene expression of the top 3 most upregulated (H) and downregulated (I) genes.

We next focused on top DEGs that were altered (Log**2**FC **;: ±**0.25) in over 70% (20/28) of the cell clusters in *Ascl1*-OE tumors. Overall, 39 top DEGs were upregulated, 14 of which were ribosomal protein genes (*Fau*, *Rpl10, Rpl12, Rpl15, Rpl23, Rpl23a, Rpl34, Rpl8, Rplp0, Rps20, Rps27a, Rps3a1, Rps4x, Rpsa*) (**Fig. 8C**), while the majority of the remaining 25 upregulated DEGs are essential for promoting cancer metastasis and invasion (*Tmsb4x, Apoe, Cst3, Sparcl1, Fabp5, Gfap, Ier2, Ntsr2, Gal*)^49–55^, NSC/GSC maintenance (*Id1, Id3*, *Fos, Hes5, Sparcl1, Sh3glb1, Lgals1*, *Gfap, Tmsb4x*)^56–61^, and/or resistance to chemotherapy (*Apoe, Mt1, Mt3, Lgals1, Sh3glb1*)^59, 62–64^ (**Fig. 8F**). Some of these top upregulated DEGs have also been reported to be the most highly expressed genes (*Apoe, Cst3, Sparcl1, Ckb, Slc1a2, Mt1, Mt3*) within cortical astrocytes across all stages of development^65^. In contrast, several of the top 25 DEGs that were downregulated are known to result in suppression of inflammatory and immune responses (*Psenen, Tbcb, Tmem147, S100b*)^66–69^, prevent NSC differentiation (*Clip3*)^70^, or promote cancer metastases (*Tbcb*)^71^ (**Fig. 8G**). The top three most upregulated DEGs are *Tmsb4x, Apoe,* and *Cst3,* which were expressed at high levels in MC3 microglial/mesenchymal cell clusters of control tumors, consistent with their roles as markers of reactive microglial^72^, but were uniformly upregulated by 2-3 fold across all cell clusters of *Ascl1*-OE tumors (**Fig. 8H**). Conversely, *Cox6b1, Psenen,* and *Tbcb* were the top three genes that were expressed at high levels in control tumor cells, but were drastically downregulated across *Ascl1*-OE tumor cells (**Fig. 8I**).

## DISCUSSION

Defining hallmarks of GBM include high degrees of inter- and intra-tumoral cellular and molecular heterogeneity combined with rapid proliferation and invasion of the brain. Studies of both human and genetically engineered mouse models implicate genomic alterations, driver mutations, cell-of-origin, and/or the tumor microenvironment as potential determining factors of the heterogeneity, lineage hierarchy, and aggressiveness of GBMs. However, precisely how these factors are translated within tumor cells to generate a variety of molecularly and behaviorally distinct GBM cellular subtypes remains unclear. In this study, we showed that in the presence of driver mutations in cells such as NPCs residing in the subventricular zone, activation of intracellular oncogenic pathways (i.e. Ras, Akt) leads to the dysregulation of ASCL1 and OLIG2, which is a critical step in the initiation of gliomagenesis. This dysregulation likely occurs at both the transcriptional and protein levels, wherein the expression of ASCL1 and/or OLIG2 further reciprocally and positively regulate each other’s expression (**Fig. S3**). In parallel, ASCL1 and OLIG2 activity may also be modulated by serine-threonine kinase driven phosphorylation resulting in stabilization of these transcription factors to promote activation of cell cycle and NSC programs^22, 73–77^. Consequently, affected NPCs are then transformed into tumor-propagating cells, which are marked by highly dynamic levels of ASCL1 and OLIG2 (**Fig. 3F,G**).

Loss- and gain-of-function studies combined with scRNA-seq reveals that tumor cells with higher levels of ASCL1 relative to OLIG2, as seen in the *Olig2*-CKO and *Ascl1*-OE tumors, favor activation of an NSC/astrocyte program (**Fig. 6E,H**), characterized by high expression of GFAP and a highly migratory or diffuse phenotype (**Fig. 4I-K,Q-S**). We propose that these ASCL1^high^ cells, which are upregulated with ribosomal protein, translation elongation factor, mitochondrial, NSC, and cancer metastasis and therapeutic resistance genes (**Fig. 8**), are the GSCs of GBM tumors. In contrast, tumor cells with higher levels of OLIG2, such as *Ascl1*-CKO tumors (**Fig. 3G**), favor activation of an OPC/oligodendrocyte program. These OLIG2^high^ cells include OPC-like cancer cells marked by SOX10 and PDGFRA (**Fig. 4E-G; Fig. S2B,F**). GSCs and OPC-like cancer cells then eventually give rise to more differentiated CNS (astrocyte, NFOL-MOL, neuronal) cancer cells, or aberrantly to non-CNS (microglial/mesenchymal, pericyte, endothelial) cancer cells within GBM tumors, in part as a result of the pioneering and reprogramming properties of ASCL1 and OLIG2^21, 78, 79^. The remnants of residual NSC and OPC gene expression within these more differentiated tumor cell-types are supportive of this interpretation (**Fig. 6I**).

ChIP-seq and co-immunoprecipitation assays for ASCL1 and OLIG2 in two orthotopic lines of GBMs reveal novel insights into the redundant and intricate binding of these two transcription factors. Notably, although OLIG2 binds more genomic sites than ASCL1, there is extensive overlap with ASCL1 binding sites. This binding overlap may be due to the direct dimerization between ASCL1 and OLIG2. Target genes associated with ASCL1 and OLIG2 shared binding, are comprised of a complex transcriptional network essential for promoting and sustaining GBM cells in a persistent state of undifferentiation, proliferation, and malignancy. Major target genes of this transcriptional network include *ASCL1* and *OLIG1/2* themselves, their respective bHLH E-protein co-binding partners, NOTCH signaling, cell cycle, and a multitude of NSC, astrocyte, and oligodendrocyte lineage genes. The significance of this transcriptional network in gliomagenesis was confirmed by their correlated expression with both *ASCL1* and *OLIG2* in RNA-seq of TCGA GBM samples. More importantly, we also validated that genes of this transcriptional network and the cell types associated with their function were altered accordingly in the absence or elevated levels of ASCL1 and/or OLIG2 in the various mouse brain tumor types (**Figs.4-8**).

Similar to gliogenesis, we showed that tdTOM+ tumor cells induced from NPCs in the subventricular zone of this GBM mouse model are predominantly restricted in fate, either as GFAP+ astroglial-like or SOX10+;PDGFRA+ oligodendroglial-like cancer cells (**Fig. 4A-C; Fig. S2A,B**). This lineage restriction was also observed in *Ascl1*-OE tumors (**Fig. 4Q-S**), where nearly all tumor cells co-expressed both ASCL1 and OLIG2 (**Fig. 3E; Fig. S3K**). This result was unexpected since these transcription factors are required developmentally for the specification and proliferation of OPCs^24, 28, 30, 80, 81^. Moreover, despite the shared binding of ASCL1 and OLIG2 to signature genes of both NSC/astroglial and OPC/oligodendroglial lineages, scRNA-seq of *Ascl1*-OE tumor cells revealed an upregulation of NSC/astrocyte signature genes but a downregulation of OPC/oligodendrocyte lineage genes (**Figs. 6**, **7**). A possible explanation for this finding is that similar to that observed in NSCs^82^, a sustained high level of ASCL1 leads to an upregulation of *Hes5* (**Fig. 5I**), which in turn downregulates OLIG2, as revealed at both the transcript and protein levels in *Ascl1*-OE compared to control and *Ascl1*-CKO tumors (**Figs. 3G**, **7A**). Consequently, this downregulation of OLIG2 results in decreases in oligodendrocyte lineage genes.

Of significant importance to the hierarchy and heterogeneity of GBMs is that the *Ascl1*-OE phenotype, combined with the *Olig2*-CKO phenotype, offers two important insights into the functional interactions between ASCL1 and OLIG2, and how they are essential for regulating opposing NSC/astrocyte-like versus OPC/oligodendrocyte-like cells in brain tumors, respectively. The first is that without OLIG2, ASCL1 cannot efficiently activate transcription of oligodendrocyte lineage genes, indicating that ASCL1’s binding to these genes may be mediated by the direct dimerization with OLIG2. Secondly, OLIG2 likely suppresses NSC/astrocyte fate in part through this direct dimerization, which may recruit ASCL1 to preferentially bind to OPC/oligodendrocyte over NSC/astrocyte lineage genes. Thus, in the absence (*Olig2*-CKO) or presence of low levels of OLIG2 (*Ascl1*-OE), ASCL1 escapes this dimerization or inhibition to potently activate NSC/astrocyte genes. The marked increase in GFAP of both *Ascl1*-OE and *Olig2*-CKO tumors compared to their respective control and dCKO tumors (**Fig. 4**) are indication that this “astrocytoma” phenotype is directly dependent on and driven by ASCL1. Consequently, the functional impact of ASCL1’s function in the context of GBM extends beyond just cell-type/subtype commitment, but also to produce highly migratory and diffuse tumor cells.

The scRNA-seq of thousands of FAC-sorted tumor cells from the GBM mouse model reveal a complex transcriptome similar to human GBM tumor cells (**Fig. 5**). Notably, all GBM subtypes (proneural, classical, mesenchymal) were represented in the mouse brain tumors, and as expected, the proneural subtype was expanded at the expense of the classical subtype with overexpression of ASCL1 due to its function as a proneural factor (**Fig. 5J**). By assigning cell types based on curated signature gene lists (**TS5)**, we found that all ten predicted cell types consisting of quiescent and activated NSC, astrocyte, OPC, NFOL, MOL, neurons, microglia, pericyte and endothelial cells were also represented within mouse brain tumors. These cell types are similar to those recently reported in primary GBMs of patients, where scRNA-seq revealed the presence of NPC or glial progenitor cancer cells at the apex of several GBM lineages of OPC/oligodendroglial, astroglial, neuronal, and mesenchymal cancer cells^18, 34^. Notably, the NPC/glial progenitor cancer cells are directly marked by ASCL1 or its direct target genes (*DLL3, HES5*), and serve as highly proliferative GSCs that are resistant to TMZ treatment^34^.

In agreement with these findings, we show that a high level of ASCL1 is a defining factor that directly confers GSCs with highly proliferative, migratory, and therapeutic resistance potential by directly or indirectly upregulating genes essential for maintaining cancer stemness and promoting chemoresistance, cancer metastases and invasion (**Fig. 8**). Notably the top upregulated gene, *Tmsb4x* encoding for Thymosin b-4, may directly mediate the migration and invasion of GSCs in the brain, as it is a potent regulator of actin polymerization^83^. Indeed, overexpression of *Tmsb4x* positively regulates NPC expansion, while silencing of *Tmsb4x* promotes stem cell differentiation^84, 85^. Similarly, downregulation of *Tmsb4x* also decreases invasion and proliferation of glioma cells, while overexpression confers stemness and chemotherapeutic resistance^84, 85^.

*Ascl1*-OE tumors also exhibited significant upregulation of genes encoding ribosomal protein small and large subunits, indicating an increased in translation and protein synthesis in these tumor cells (**Fig. 8B,C**). These biological processes are similarly increased in cancer stem cells (CSCs). More specifically, CSCs or proliferating progenitor cells exhibit elevated expression of ribosomal protein subunit genes, while these genes are decreased in post-mitotic cells and following cell-fate commitment^43, 86, 87^. We found that nearly 90% (71 of 80) of ribosomal protein subunit genes were upregulated in *Ascl1*-OE tumor cells, including those associated with chemoresistance (*Rps13*), stemness (*Rpl10a*), immunosuppression (*Rps19*), proliferation (*Rps13, Rpl32, Rplp2, Rpl35*), migration/invasion (*Rpl27a, Rplp1, Rplp2, Rpsa, Rpl35*), and poor prognosis in solid tumors (*Rps11, Rps20, Rpl30*)^88–101^.

Finally, GSCs require metabolic versatility to survive in hypoxic microenvironments and to provide energy for metabolically costly processes such as migration and proliferation^102^. CSCs are known to exhibit significant dependence on oxidative phosphorylation, which is more efficient in producing ATP than glycolysis^103^. We saw significant upregulation of both nuclear (*Cox4i1, Cox5a, Cox5b, Cox6c, Cox7a, Cox7c, Cox8a*) and mitochondrial (*mt-Co1, mt-Co2, mt-Co3*) genes encoding for the 13 subunits of cytochrome c oxidase in the electron transport chain, the key regulator of oxidative phosphorylation^104, 105^. More specifically, increases in COX4I1 isoforms are implicated in poorer prognosis, supercomplex formation, increased tumor growth and metastases, and radioresistance^106–108^. Therefore, through upregulation of cytochrome c oxidase subunits, especially *Cox4i1*, ASCL1 increases the efficiency of oxidative phosphorylation and, ultimately, confers a GSC phenotype.

In summary, the presence of multiple GBM subtypes and cell types in brain tumors of our mouse model developed and reported here highlights the robustness of our study to recapitulate the cellular and molecular dynamics of human GBMs. Our comprehensive *in vivo* analyses of ASCL1 and OLIG2 loss- and gain-of-functions demonstrate that these two bHLH transcription factors operate at the apex of gliomagenesis to intrinsically determine the hierarchy and heterogeneity of GBM cells. Importantly, the various glioma tumor types generated in the brains of this immunocompetent mouse model combined with identification of GSC signature genes upregulated by a high level of ASCL1 offer new opportunities to further elucidate and mitigate the mechanism of therapeutic resistance for GBMs.

## Acknowledgments

This research was supported by NINDS K22NS09267 & R01NS121660 (T.Y.V.), NIAAA T32 AA 014127 Alcohol Research Training in Neuroscience grant (B.L.M.), Cancer Prevention and Research Institute of Texas RP130464 (J.E.J), and partially supported by the UNM Comprehensive Cancer Center Support Grant NCI P30CA118100, ACS-IRG#131567, Pilot Projects #1451,1513, and the University of Colorado Anschutz Medical Campus Genomics Shared Resource Cancer Center Support Grant P30CA046934. The following UNM Comprehensive Cancer Center Shared Resource facilities, which receives additional support from the State of New Mexico, were essential toward the completion of this research: Analytical/Translational Genomics, Fluorescence Microscopy, Flow Cytometry, Bioinformatics, and Human Tissue Repository. *FUGW-Cre* plasmids were a gift from Dr. Jason Weick’s lab at UNM. We thank Drs. Nora Perrone-Bizzozero and Fernando Valenzuela for critical reading of the manuscript.

## Author Contributions

BLM and TYV designed the study, and BLM carried out all mouse brain tumor experiments. KJB and BLM performed bioinformatic analyses of scRNA-seq data. LEPB, MSK, JN, RA, YL, and CMM harvested brain tumors and performed immunohistochemistry. HS performed Co-IP experiments and characterized plasmid DNA. TYV and MDB performed ChIP-seq and RK performed bioinformatic analyses of ChIP-seq and TCGA GBM RNA-seq under supervision of JEJ, who also contributed to early stage of this study. RMB provided PDX-GBM mice. BLM and TYV analyzed all data, prepared figures, and wrote the manuscript. All co-authors provided edits and approved the final manuscript.

## MATERIALS and METHODS

### ChIP-seq and TCGA RNA-seq data analyses

Details of our ChIP-seq assays for ASCL1 (GSE152401) using two orthotopic lines of patient-derived GBM xenograft (PDX-GBMs) (R548 and R738) were previously reported^22^. Here, ChIP-seq assays for OLIG2 were done for the same two PDX-GBM lines to directly compare binding profiles of ASCL1 and OLIG2. Briefly, PDX-GBMs were grown and dissected from brains of NOD-SCID mice and then homogenized and fixed in 1% formaldehyde to crosslink proteins and DNA, followed by quenching with 0.125 M of glycine. Nuclear chromatin was pelleted, washed with cold PBS, and sonicated into 200–300 bp fragments using a Biorupter (Diagenode). A portion of the sheared chromatin (10%) was set aside as input DNA, while 100 μg was subjected to chromatin immunoprecipitation (ChIP) using 5 μg of mouse anti-ASCL1 (Mash1) antibody (BD Biosciences, 556604) or 5 μg rabbit ant-OLIG2 (Olig2) (Millipore, AB9610). Washes and reverse-crosslinking were performed using Dynabeads Protein G to elute the ChIP DNA for sequencing.

ChIP-seq analyses were performed as we previously reported^22^. Briefly, bowtie2 (v2.2.6)^109^ was used to aligned sequence reads to the human reference genome (hg19) and *de novo* motif analyses were performed using HOMER Software (v.4.7). Target genes associated with ASCL1 or OLIG2 ChIP-seq peaks were determined using GREAT v4.0.4 (http://great.stanford.edu/public/html/)^35^. We used Spearman rank order correlation (>0.4) to identify the top 10% of genes found in bulk RNA-seq of GBM samples in the TCGA cohort^13^ positively correlated with *ASCL1* and/or *OLIG2* expression^22^. Biovenn (https://www.biovenn.nl/) was used to generate Venn diagrams of overlapped ASCL1 and OLIG2 target genes identified from ChIP-seq with *ASCL1* and *OLIG2* positively or negatively correlated identified from TCGA GBM RNA-seq.

### Co-immunoprecipitation

PDX-GBM tumors or brains of E14.5 mouse embryos were carefully dissected and placed in cold PBS with protease inhibitors. Tumor or brain tissues were then chopped and transferred to glass tubes containing 37°C DMEM for homogenization with glass dounce tissue grinders. Cells were then washed with cold PBS, resuspended, and lysed in RIPA buffer to isolate supernatant of nuclear extracts, which were then pre-cleared with Dynabeads Protein G. Immunoprecipitation was performed by adding anti-ASCL1/Mash1 antibody (BD Biosciences, 556604) for 2 hours followed by addition of Dynabeads Protein G rotated at 4°C overnight. Following incubation at 95°C for 5 minutes, both immunoprecipitation products and input proteins were extracted and subjected to Western blot electrophoresis (LI-COR Quick Western Kit) for detection using mouse anti-ASCL1 and anti-rabbit OLIG2 (Millipore/AB9610) on nitrocellulose followed by the appropriate Alexa680/IR800 secondary antibodies. All Western blots were imaged on an Odyssey scanner (LI-COR Biosciences).

### Mouse strains used for this study

All mouse experiments in this study followed NIH guidelines and a research protocol was approved by the Institutional Animal Care and Use Committee (IACUC) at the University of New Mexico Health Sciences Center. The transgenic mouse lines used for this study and their sources are shown in Supplementary Table 1 (**TS1**).

### Inducing brain tumors in transgenic mice by electroporation of Cre and CRISPR-Cas 9+gRNA plasmids

Transgenic mouse lines were crossed to carry a combination of floxed and reporter alleles to generate control, *Ascl1*-conditional knock-out (CKO), *Olig2*-CKO, *Ascl1; Olig2*-double CKO; or *Ascl1*-overexpression (OE) tumors (**Table 1**). Brain tumors were induced in mouse pups on the day of birth (P0) by injection of approximately 1 μL volume of the following plasmid mix and concentrations (*FUGW-Cre* [2 μg/μL] + *pX330-Cas9+gNf1* [1 μg/μL] + *pX330-Cas9+gPten* [1 μg/μL] + *pX330-Cas9+gTp53* [1 μg/μL]) into the lateral ventricle, followed by electroporation into NPCs/GPs lining the right dorsal subventricular zone (**Fig. S1**). The guide RNAs used were designed to target the following sequences of each of the three tumor suppressor genes^37^: *Tp53* 5’-ACAGCCATCACCTCACTGCA-3’, *Pten* 5’-AAAGACTTGAAGGTGTATAC-3’, and *Nf1* 5’-AGTCAGCACCGAGCACAACA-3’. Electroporation was performed using a NEPA21 Super Electroporator and CUY650-P5 electrokinetic tweezers with 5 mm platinum disk electrodes placed diagonally directly above the right lateral ventricle (+ disk) and below the jaw (- disk) of mouse pups (**Fig. 2A**). Each pup underwent electroporation twice with 5 pulses (100V, 50 ms duration, 950 ms interval) about 5 minutes apart to ensure efficient force transfer of plasmids into cells in the subventricular zone and 100% tumor penetrance. A 2:1 concentration ratio of *Cre* to *gRNA* plasmids was used to ensure that tumors were labeled by tdTOM or GFP in the dorsal cortex.

### Brain tumor harvest and survival curves

Brain tumors were harvested from mice at P30, P60, or if severe neurological symptoms (hunching, seizures, etc.) were observed; these symptoms were considered endpoints for survival curve and terminal tumor analyses. We verified that neurological symptoms were due to a brain tumor mass labeled by fluorescent reporter in the right hemisphere, rather than other non-tumor related complications (i.e. hydrocephalus) which can occur in some mice. Longitudinal analyses and survival curves were based on data from both male and female tumor-bearing mice from at least 3 litters per genotype. Statistical significance and median survival were determined by simple survival analysis (Kaplan-Meier, Prism 10) between control versus each experimental tumor group, or between male versus female of the same tumor group.

### EdU injection and immunohistochemistry of brain tumor sections

Two hours before tumor harvest, 5-ethynyl-2′-deoxyuridine (EdU, 1 mg/mL dissolved in sterile PBS), a thymidine analog was injected (10 μg EdU/g body weight) intraperitoneally to label proliferating tumor cells. To harvest brain tumors, mice were intraperitoneally injected with Avertin (2.5 g 2,2,2 tribromoethanol [Aldrich T4,840-2] + 5 mL 2-methy-2-butanol (amylene hydrate [Aldrich 24,048-6] + 200 mL distilled water) to induce anesthesia. Hearts were then perfused with 4% PFA/1XPBS. Brains were extracted and placed in 4% PFA/1XPBS overnight, washed in 1XPBS followed by submersion in 30% sucrose/1XPBS, and then frozen embedded in O.C.T. compound for cryosectioning.

For immunohistochemistry, brains were cryosectioned at 40 μm thickness and then blocked as floating sections for one hour with 2% Goat/Donkey blocking solution with 0.3% Triton X-100, followed by overnight incubation in primary antibodies diluted in the blocking solution. The next day, the floating sections were incubated in the appropriate secondary antibodies conjugated to Alexa fluorophores (488, 568, or 647; ThermoFisher). For EdU staining, brain tumor sections were incubated for 30 minutes in detection solution containing 100 mM sodium ascorbate, 4 mM copper sulfate, and 12 μM Cy5 in PBS as previously described^110^. The primary antibodies used in this study are listed in Supplemental Table 1 (**TS1**). Hematoxylin and eosin (H&E) staining of mouse brain tumors was performed by the Human Tissue Repository core at UNM.

### Quantification of tumor cells positive for EdU, ASCL1, or OLIG2

Brain sections were imaged using confocal microscopy (Leica TCS SP8). Images were collected using the 20X oil objective at 2048×2048 resolution (individual image size: 581 μm x 581 μm). Imaging was limited to regions within tumors or on tumor margins with the highest density of EdU and reporter (tdTOM or GFP) co-labeled cells. EdU+/reporter+, ASCL1+/reporter+, and OLIG2+/reporter+ staining was quantified with IMARIS software. In brief, 581 μm x 581 μm images with an optical density of 3.12 μm were converted from TIFFs to .ims files and a colocalization channel was created using DAPI and the tumor reporter to limit quantification to DAPI+/reporter+ tumor cells. Using the Spots application, we assessed the number of nuclei with an estimated average diameter of 6.8 μm within the colocalization channel. Next, another channel (EdU, ASCL1, or OLIG2) was added to the Spots application and assessed again using “Classify Spots”. This analysis quantified cells that were positive for the reporter and the channels of interest. We analyzed 5 images of distinct tumor areas per coronal brain slice and 3 brain slices per tumor, for a total of 15 images per brain at terminal stages. Fewer images analyzed for P30 brains because of smaller tumor sizes, but at least 5 images per brain were analyzed.

### Quantification of cellular immunofluorescent levels of ASCL1 and OLIG2 within tumor cells

Sections of control, *Ascl1*-CKO*, Ascl1*-OE, and *Olig2*-CKO brain tumors were simultaneously stained with the same anti-ASCL1 or anti-OLIG2 antibody concentrations and imaged using the same confocal laser parameters. Cellular immunofluorescence intensity for ASCL1 or OLIG2 was calculated as previously described^80, 111^. Briefly, single tumor cells within high magnification images (TIFF) with ASCL1 or OLIG2 signals were encircled using the freeform drawing tool in ImageJ software. The area, integrated density, and mean fluorescence of individual cells, along with mean background fluorescence, were measured for calculation of the total corrected cellular fluorescence (TCCF=integrated density – (area of cell x mean background fluorescence)). We measured TCCF for 20 cells per tumor section and 3 sections per tumor (60 cells total).

### Quantification of tumor cell migration distance across the midline on contralateral corpus callosum

Tile scans of whole coronal brain sections with visible corpus callosum and labeled tumor cells were collected for each brain at three different rostral-caudal levels through the bulk of the tumor. Using ImageJ software, we captured the total length (TL) of the corpus callosum on the contralateral hemisphere starting at the midline, followed by the migration distance (MD) of tumor (reporter+) cells on the contralateral corpus callosum. The distance of tumor migration on the contralateral corpus callosum was then determined by dividing MD/TL, and reported as percentage (%) distance migration on the contralateral corpus callosum for each tumor sample. This normalization was used because large tumor masses, such as those in control and *Ascl1*-CKO tumor mice, can drastically alter the size, absolute length, and morphology of the contralateral corpus callosum.

### Single cell suspension and fluorescence-activated cell sorting (FACS) of tumors

Brains of control (tdTOM+, N=3) or *Ascl1*-OE (GFP+, N=3) tumor-bearing mice with neurological symptoms were freshly harvested and tumors were dissected out in a bath of Hank’s Balanced Salt Solution (without calcium or magnesium). Working on ice, tumor tissue was weighed and then transferred to a gentleMACS C tube and dissociated using the Miltenyi Neural Tissue Dissociation Kit (P) and an Octo-Dissociator with Heaters. Following this, cells underwent FACS in the UNM Comprehensive Cancer Center Shared Resource Flow Cytometry facility. Tumor cell concentrations and viability were determined using a Cell Countess II FL device (ThermoFisher) before calculating cell counts and loading the suspension into the Next GEM Chip G and Chromium Controller (10x Genomics) per the manufacturer’s protocol for genome-scale metabolic models (GEM).

### Library preparation of FAC-sorted tumor cells for 10X Genomics Chromium scRNA-seq

Tumor cell concentrations and viability were determined using the Cell Countess II FL (ThermoFisher) prior to calculating cell counts and loading the suspension into the Next GEM Chip G and Chromium Controller (10x Genomics) per the manufacturer’s protocol for GEM formation. The 10x Chromium Next GEM 3’ protocol was used to create 3’ libraries for sequencing at the University of Colorado Anschutz Medical Campus’s Genomics Shared Resource Cancer Center using Illumina NovaSEQ 6000 instruments on S4 flow cells. Briefly, cells were lysed and barcoded within each GEM before first strand cDNA synthesis, and also within each GEM. Cells were pooled prior to library completion as described in the manufacturer’s protocol. Library quality was assessed after cDNA synthesis and after completion on the BioAnalyzer using a DNA High Sensitivity Chip (Agilent). Before sequencing, Agilent Tape Station 4200 and Invitrogen Qubit 4.0 reagents were used to determine final library concentrations before dilution, normalization, and pooling at 4 nM. qPCR was used to determine cluster efficiency before loading libraries into NovaSEQ devices.

### Single-cell RNA sequencing analyses of tumor cells

Data were demultiplexed and fastq files were generated using bcl2fastq (v2.20.0.422, Illumina) with parameter “--barcode-mismatches” set to 1. Fastq files were aligned and genes/cells were counted against the mouse reference genome (mm10) using CellRanger (v6.0.0, 10xGenomics)^112^. EGFP and tdTomato sequences were appended to the mm10 genome according to the manufacturer’s instructions. Approximately 18,000 tdTOM+ control tumor cells and 25,000 GFP+ *Ascl1*-OE tumor cells met quality criteria for further analyses. CellRanger-generated filtered files for these cells were used for downstream analyses.

Seurat (v4.3.0.9002), an R package (v4.3), was used for downstream unsupervised clustering analyses (https://www.R-project.org/)^112, 113^. We used scatter (v1.26.1) to identify and remove cells that were outliers for counts, features, and mitochondrial counts^114^. scDblFinder (v1.12.0) was used to identify and remove doublet cells^115^. Data were transformed using SCTransform (v0.3.5)^116^ as implemented in Seurat. Data from all samples were integrated using Seurat’s standard integration workflow. Principal component analyses and an elbow plot were used to visualize variances and select principal components (1:40). Clusters were determined using the FindNeighbors and FindClusters function with default parameters and the resolution set to 1.1. GBM subtype annotations were added using Seurat’s MetaFeature function and a list of subtype markers previously described^13^. Cell cycle phases were determined using the CellCycleScoring function within Seurat with a list of mouse cell cycle genes from https://hbctraining.github.io/scRNA-seq/lessons/cell_cycle_scoring.html. Cell type prediction was performed using singleR (v2.0.0) based on a curated list of cell type-specific signature genes (**Table S5**) adopted from previously published datasets (brainrnaseq.org)^40–45^. Differential gene expression was determined using FindConservedMarkers (difference between clusters/cell types) or FindAllMarkers (differences between control and treated for each cell cluster/cell type) in conjunction with the R package MAST (v1.24.1)^117^ as implemented in Seurat. Significant genes were defined by average Log2 fold-change >±0.25, adjusted p-value < 0.05, and expressed in > 10% of the cells in the cluster. Unionized pseudobulk RNA-seq results for transcripts of each cell type were calculated by AverageExpression within Seurat with the grouping for each cell type for each tumor. All heatmaps of the pseudobulk RNA-seq were generated using the R package ComplexHeatmap (v2.14.0)^118, 119^, wherein columns were ordered manually by tumor ID and then cell type annotation and rows were clustered automatically.

---

**Supplementary Figure 1.**
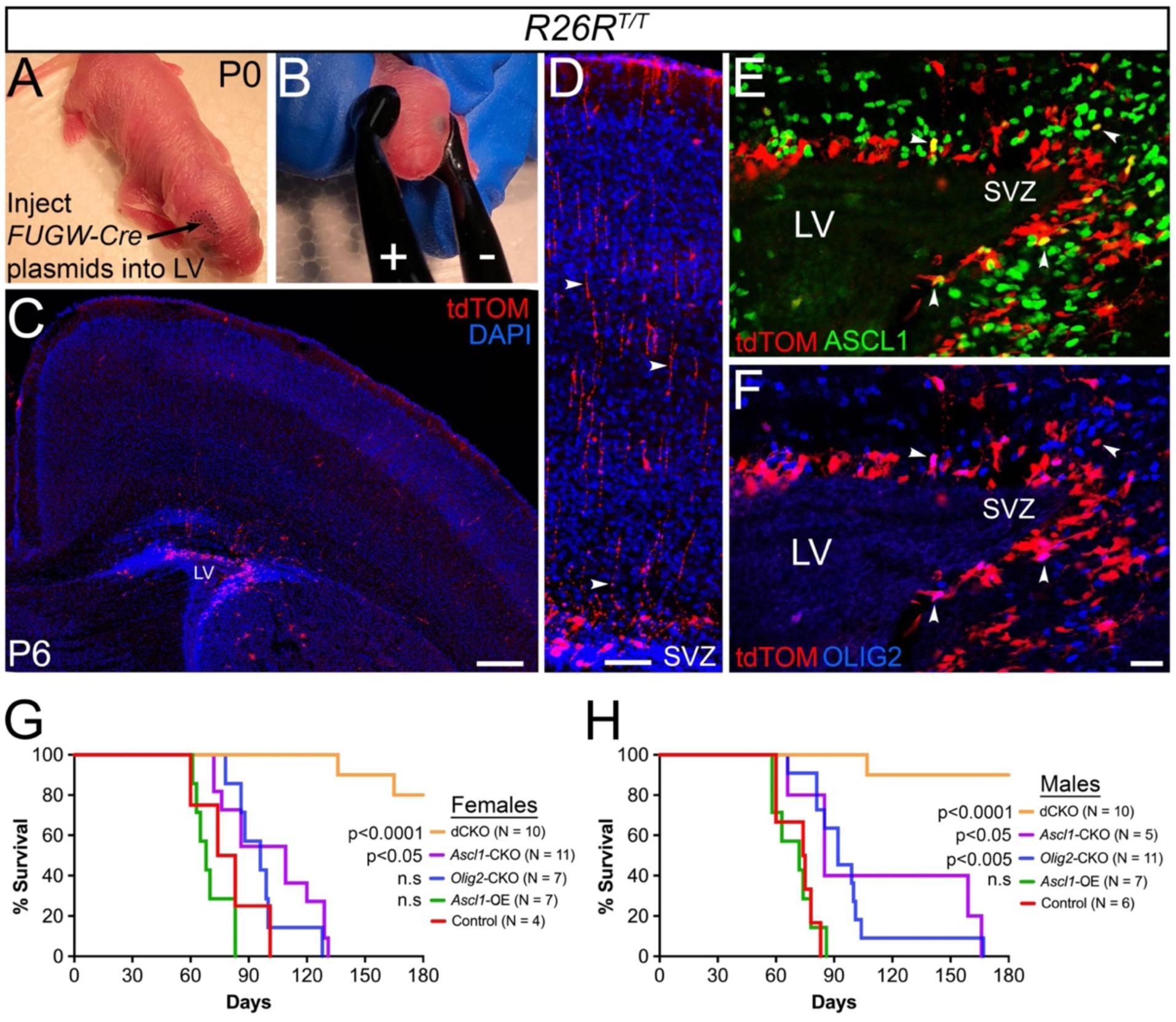
Electroporation of NPCs in the subventricular zone of *R26R^T/T^* mice. (**A,B**) Mouse pups are injected with *FUGW-Cre* plasmids into lateral ventricle and electroporated at P0. (**C,D**) tdTOM and DAPI staining at P6 showing Cre-mediated labeling of NPCs in the dorsal subventricular zone (SVZ) with radial processes extending to the pia surface (arrowheads). (**E,F**) Only a few tdTOM+ NPCs express ASCL1 and OLIG2 (arrowheads). (**G,H**) Survival curve of female and male mice for the various tumor groups. Note that there is no significant difference in survival between male and female of the same tumor type, but only between control and the indicated experimental tumor groups (Kaplan-Meier survival analysis). Scale bars: 250 μm for panel C, 100 μm for panel D, and 25 μm for panels E,F.

**Supplementary Figure 2.**
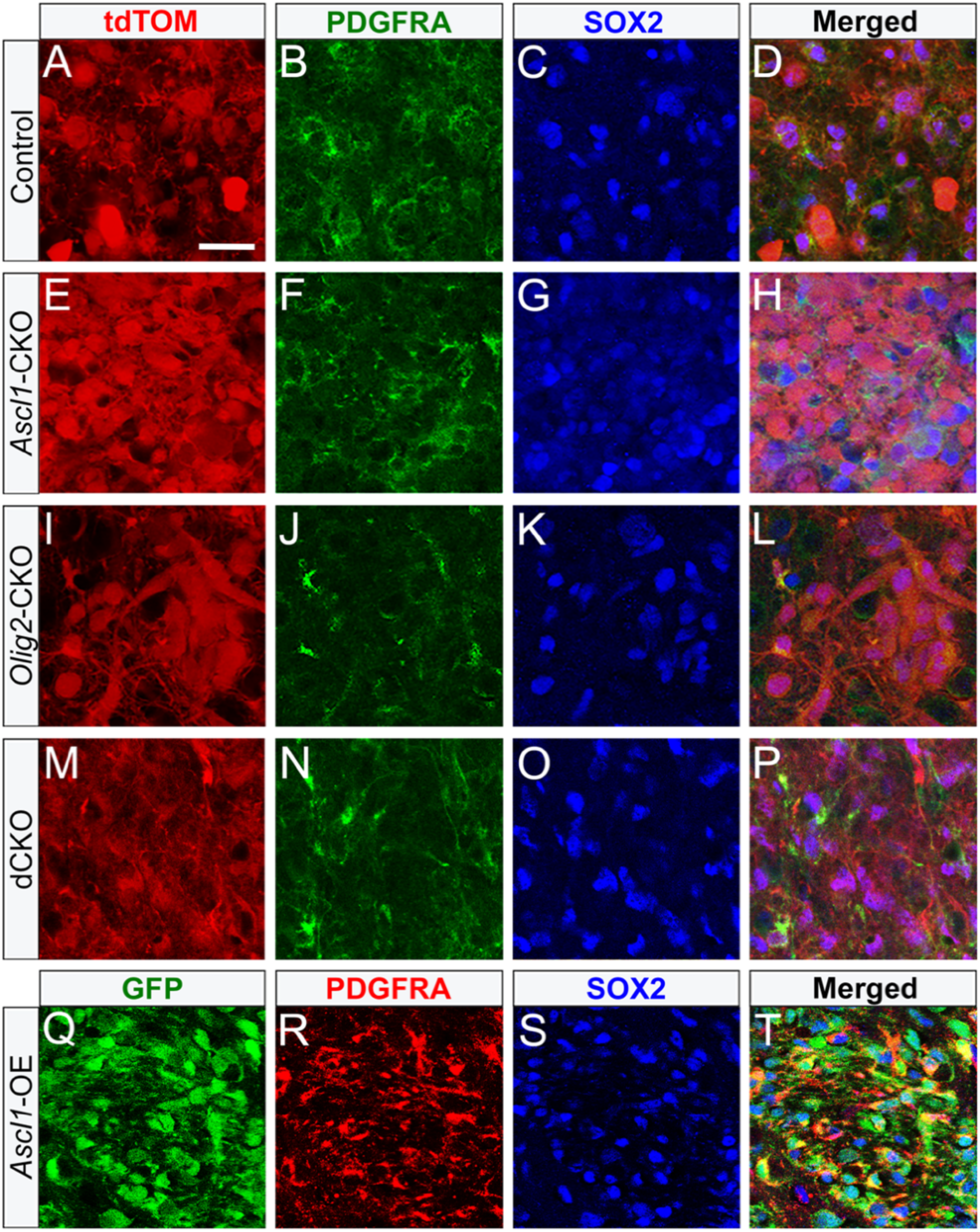
SOX2 and PDGFRA are differentially expressed in mouse brain tumors. (**A-T**) Double immunofluorescence showing that SOX2 is expressed in tumor cells of all tumor types whereas PDGFRA expression is high in control (A-D), *Ascl1*-CKO (E-H), and *Ascl1*-OE (Q-T) tumors, but low in *Olig2*-CKO (I-L) and dCKO (M-P) tumors. Scale bar: 25 μm for all images.

**Supplementary Figure 3.**
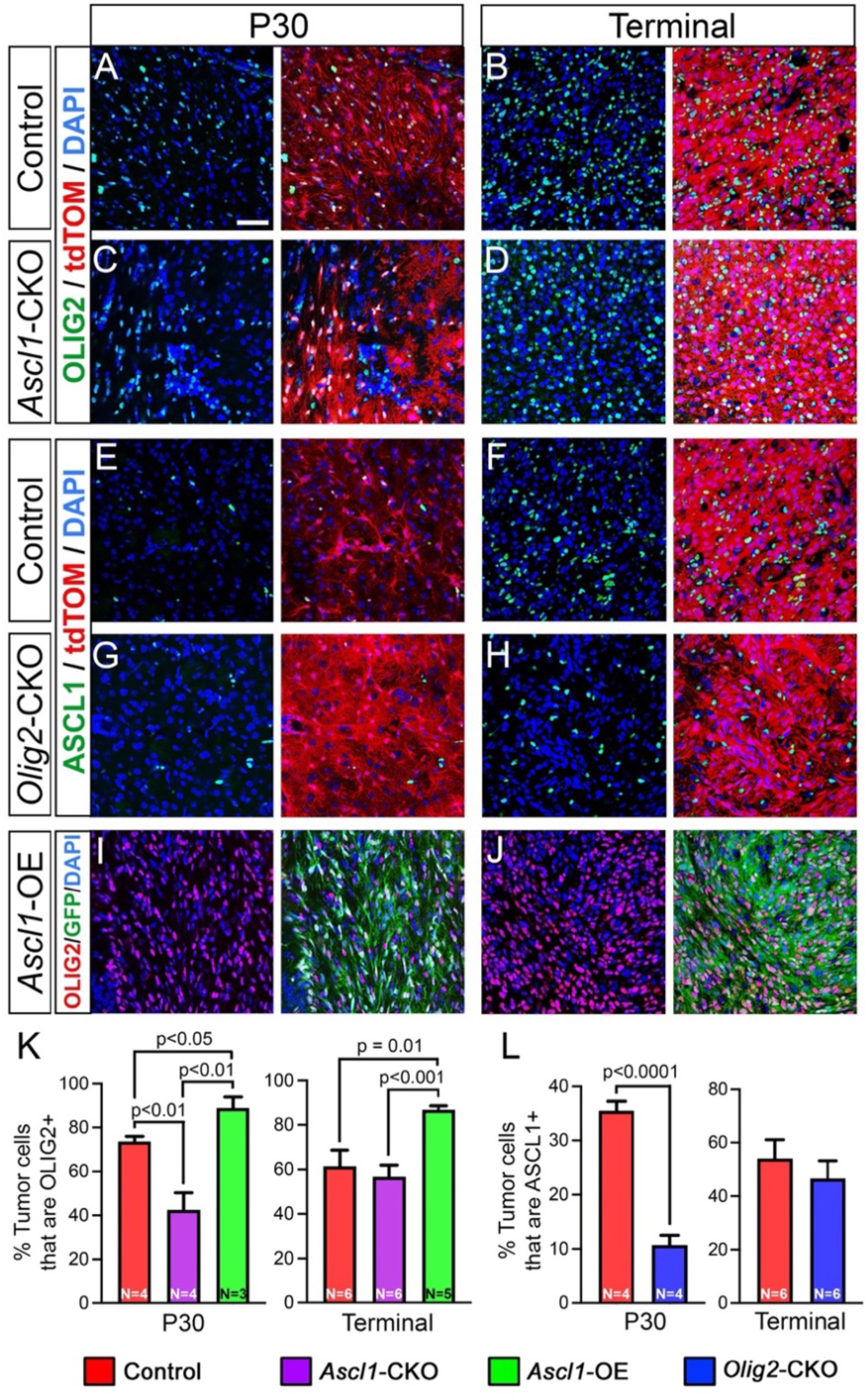
ASCL1 and OLIG2 reciprocally regulate expression in glioma tumors. (A-J) Representative immunofluorescent images of OLIG2+ tumor cells in control (A,B), *Ascl1*-CKO (C,D), and *Ascl1*-OE (I,J) tumors, or ASCL1+ tumor cells in control (E,F) and *Olig2*-CKO (G,H) tumors at P30 and terminal stages. **(K,L)** Percentage of labeled tumor cells positive for OLIG2 or ASCL1 for the different tumor types. Data shown are mean ± SEM. Statistical significance was determined using unpaired t-tests. Scale bar: 50 μm for all images.

**Supplementary Figure 4.**
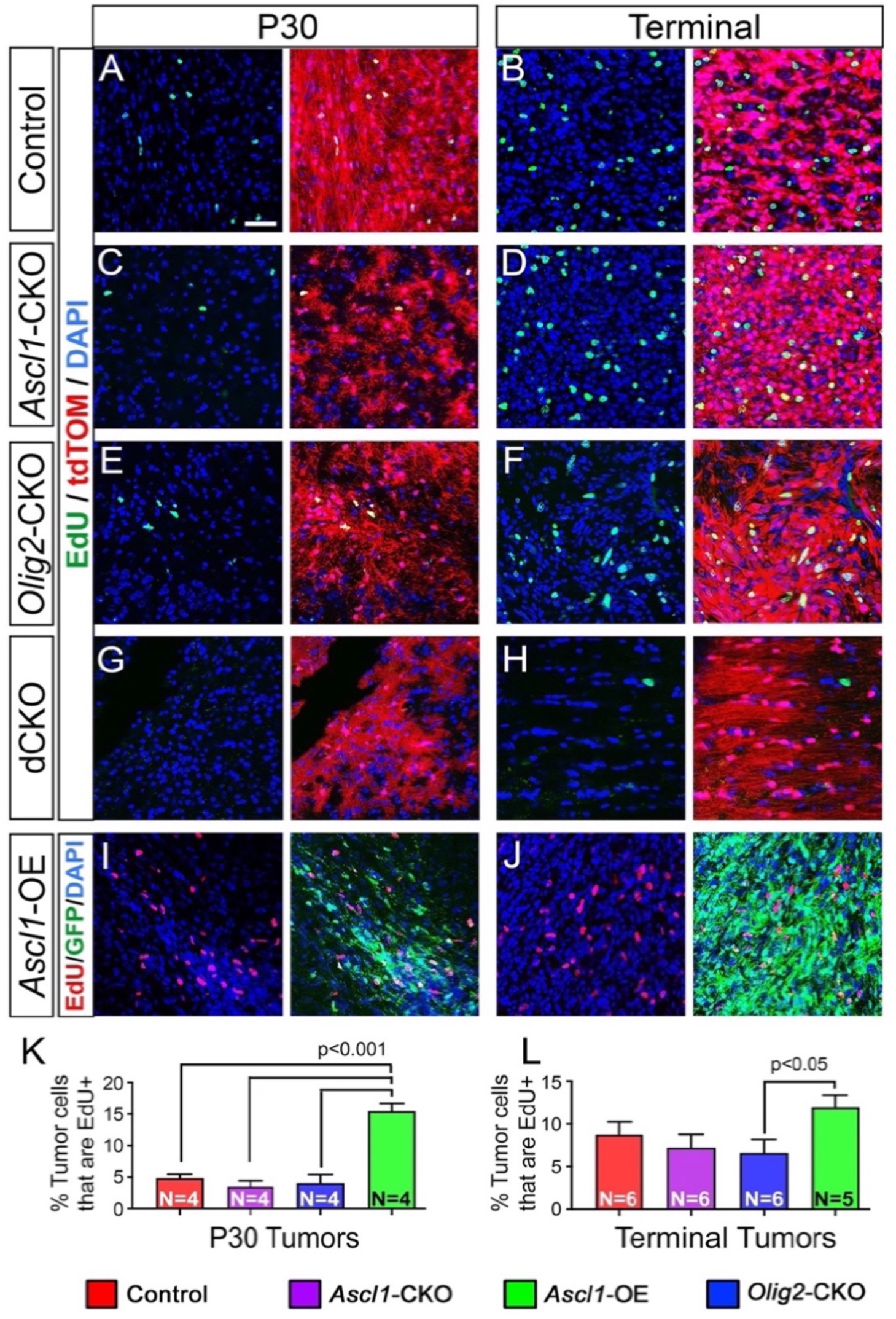
**ASCL1 and OLIG2 are required for proliferation of brain tumors**. **(A-J)** Representative images of EdU staining for control (A,B), *Ascl1*-CKO (C,D), *Olig2*-CKO (E,F), dCKO (G,H), and *Ascl1*-OE (I,J) tumors at P30 and terminal stages. **(K,L)** Percentage of tumor cells positive for EdU at P30 and terminal stages. Data shown are mean ± SEM. Statistical significance was determined using unpaired t-tests. Scale bar: 50 μm for all images.

**Supplementary Figure 5.**
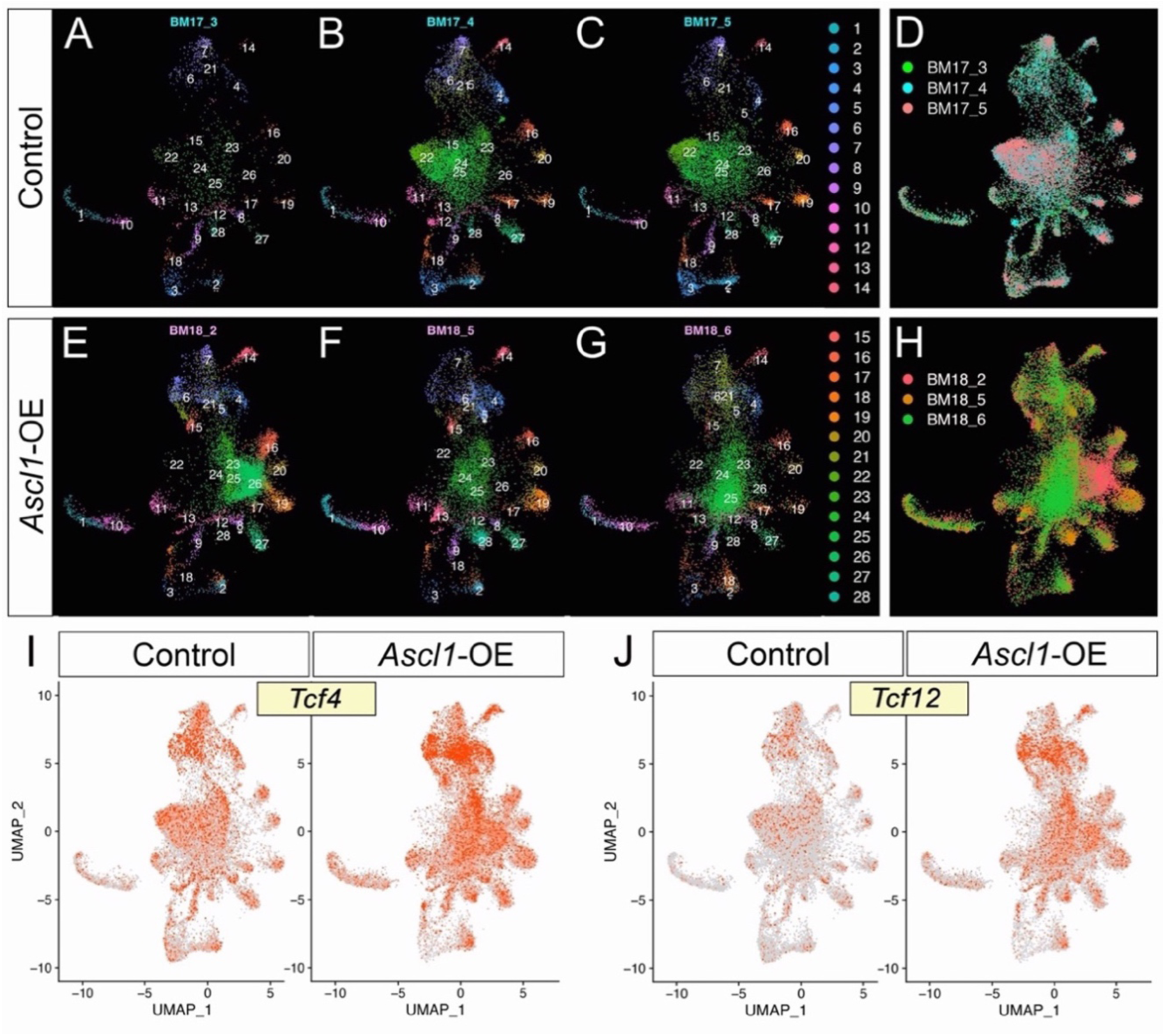
**Unsupervised clustering of cells from individual tumors**. **(A-H)** UMAP showing clustering distribution of tumor cells from each tumor sample according to genotype. Cells from each of the 3 control and 3 *Ascl1*-OE tumors are distributed across the 28 cell clusters. **(I,J)** UMAP showing increased expression of bHLH E-protein encoded genes, *Tcf4* and *Tcf12*, in *Ascl1*-OE tumor cells.

